# Stochastic resonance and bifurcations in a heterogeneous neuronal population explain intrinsic oscillatory patterns in entorhinal cortical stellate cells

**DOI:** 10.1101/2022.01.23.477388

**Authors:** Divyansh Mittal, Rishikesh Narayanan

## Abstract

Stellate cells in the medial entorhinal cortex manifest peri-threshold oscillatory patterns in their membrane voltage. Although different theoretical frameworks have been proposed to explain these activity patterns, these frameworks do not jointly account for heterogeneities in intrinsic properties of stellate cells and stochasticity in ion-channel and synaptic physiology. In this study, we first performed *in vitro* patch-clamp electrophysiological recordings from rat MEC stellate cells and found pronounced cell-to-cell variability in their characteristic physiological properties. We demonstrate that stochasticity introduced into either a simple nonlinear dynamical system (Hopf bifurcation) or into heterogeneous biophysical models yielded activity patterns that were qualitatively similar to peri-threshold oscillations in stellate cells. We developed five quantitative metrics for identification of valid oscillatory traces and confirmed that these metrics reliably captured the variable amplitude and non-rhythmic oscillatory patterns observed in our electrophysiological recordings. We analyzed traces from a stochastic Hopf bifurcation system for further confirmation on the reliability of these quantitative metrics in detecting oscillatory patterns. Apart from providing confirmation, these analyses provided a key insight about the manifestation of stochastic resonance in the stochastic bifurcation system, but not with theta-filtered noise. We then validated peri-threshold activity patterns obtained from a heterogeneous population of stellate cell models, with each model assessed with multiple trials of different levels and forms of noise (ion-channel, synaptic, and additive) at different membrane depolarizations. Strikingly, the validation process confirmed the manifestation of heterogeneous stochastic bifurcations across all models and revealed the detection of the highest number of valid oscillatory traces at an optimal level of each form of noise. The manifestation of stochastic resonance in this heterogeneous model population explained why intrinsic theta oscillations might not be observed under *in vivo* conditions where noise levels are typically high. Together, we provide several lines of evidence that demonstrate heterogeneous stochastic bifurcations as a unifying framework that fully explains peri-threshold oscillatory patterns in stellate cells and argue for state-dependence in the manifestation of these oscillations.

## INTRODUCTION

Neurons in the entorhinal cortex are positioned at a crucial stage of information processing. They receive polymodal sensory information and provide spatial information to hippocampus (Moser et al., 2017; Nilssen et al., 2019; Tukker et al., 2021). One of the spatially selective cell types in the medial entorhinal cortex (MEC) are the layer II stellates cells. Stellate cells that elicit action potentials forming a hexagonal or triangular grid-like repetitive pattern tiling the spatial environment are called as grid cells (Fyhn et al., 2004; Hafting et al., 2005; Couey et al., 2013; Schmidt-Hieber and Hausser, 2013; Gu et al., 2018). One of the signature intrinsic properties of stellate cells is the emergence of subthreshold and mixed-mode oscillations upon depolarization closer to spiking threshold (Alonso and Llinas, 1989; Alonso and Klink, 1993; Klink and Alonso, 1993; White et al., 1993; Dickson et al., 2000b; Fransen et al., 2004; Boehlen et al., 2013).

Classically, these peri-threshold activity patterns were considered to be a result of interactions between hyperpolarization-activated cyclic nucleotide-gated (HCN) and persistent sodium (NaP) ion channels giving rise to a periodic oscillator (Hutcheon and Yarom, 2000; Fransen et al., 2004). Other studies suggest that these peri-threshold activity patterns to be attributable to theta-filtered noise (Dodson et al., 2011). However, these frameworks either do not account for the pronounced cell-to-cell variability in ion-channel and intrinsic properties of MEC stellate cells or for the widespread stochasticity that spans ion-channel and synaptic physiology. In this study, accounting for heterogeneities and stochasticity, we argue that neither the periodic oscillator nor the theta-filtered noise frameworks can fully explain peri-threshold oscillations in MEC stellate cells. Instead, with several lines of evidence, we argue that these peri-threshold activity patterns and their characteristic features are fully explained by stochastic bifurcations in a heterogeneous neuronal population.

First, we performed *in vitro* patch-clamp electrophysiological recordings from rat MEC stellate cells and found pronounced cell-to-cell variability in characteristic physiological measurements. These recordings showed characteristic oscillatory patterns with variable amplitude and frequency at disparate peri-threshold voltage ranges, with spike clustering observed in mixed-mode oscillations. Heterogeneities in electrophysiological recordings emphasized the need to employ a heterogeneous population of neuronal models to assess the theoretical underpinnings behind these oscillatory patterns. Specifically, studying a single hand-tuned model with one set of active and passive properties would not capture the pronounced heterogeneities in the electrophysiological and biophysical characteristics of MEC stellate cells. This is especially the case because biological neurons, including MEC stellate cells, manifest ion-channel degeneracy, whereby disparate combinations of structural components could manifest similar characteristic physiological properties (Mittal and Narayanan, 2018; Rathour and Narayanan, 2019; Goaillard and Marder, 2021). Therefore, we employed a previously generated (Mittal and Narayanan, 2018) heterogeneous population of 155 models that were validated against characteristic electrophysiological properties of MEC stellate cells. This population of models manifested degeneracy in satisfying the characteristic physiological properties, thereby implying pronounced heterogeneities in the underlying biophysical parameters as well (Mittal and Narayanan, 2018).

A bifurcation in a dynamical system is defined as an abrupt qualitative change in the topology or phase portrait in response to small smooth change in the value of a bifurcation parameter (Strogatz, 2014). Stellate cell responses to depolarizing current injection with continually increasing amplitude vary from resting conditions to subthreshold oscillations to mixed-model oscillations to continuous firing to depolarization-induced block. Each of these transitions could be considered as bifurcations, and neurons are often modeled as dynamical systems manifesting bifurcations (Ermentrout and Terman, 2010; Izhikevich, 2010). From a mechanistic standpoint, neuronal bifurcations are mediated by the gating properties and kinetics of the specific set of ion channels expressed in each neuron. As ion channels are fundamentally stochastic in their function and neurons typically receive non-deterministic synaptic inputs (Faisal et al., 2008), it is critical that neuronal bifurcations are not treated as deterministic bifurcations, but as stochastic bifurcations. The manifestation of stochasticity implies considerable trial-to-trial variability in responses, resulting in considerable noise in the amplitude and frequency of patterns emanating from limit cycles in these dynamical systems.

To assess the impact of stochasticity on the emergence of valid oscillatory patterns, we introduced three different forms of noise (ion channel, synaptic, and additive) into the heterogeneous population of LII MEC cells. We observed the manifestation of heterogeneous stochastic bifurcations across the population of MEC stellate cell models, qualitatively demonstrating that noise could play a stabilizing role in yielding intrinsic oscillatory patterns that were similar to their physiological counterparts. To quantify such beneficiary roles of noise, we developed five different quantitative metrics, derived from the spectrogram of activity patterns, to detect the presence of stable oscillatory traces. We set bounds on these measurements such that they were sufficient to capture the variable amplitude and irregular oscillatory patterns observed in rat MEC stellate cells. We further validated the spectrogram-based metrics in capturing the stability of the oscillations in a stochastic nonlinear dynamical system (Hopf bifurcation). Recruiting the stochastic Hopf bifurcation for assessing the reliability of these measurements also provided a key quantitative insight about stochastic bifurcations: the manifestation of stochastic resonance (McDonnell and Abbott, 2009) in this stochastic bifurcation system. Specifically, our analyses of the stochastic Hopf bifurcation unveiled the presence of an optimal level of noise that facilitates stabilization of oscillatory patterns, with any other noise level (higher or lower) hampering stabilization. In striking contrast, there was no stochastic resonance expressed with increasing noise levels in theta-filtered noise traces, and thus provided a quantitative handle to assess activity patterns in MEC stellate cells.

We performed this validation process on traces from the stochastic, heterogeneous population of MEC stellate cells and found that noise-induced stabilization of peri-threshold oscillations occurred at an optimal level of noise. This expression of stochastic resonance provided a further line of evidence that activity patterns in MEC stellate cells were consistent with stochastic bifurcations, and not with theta-filtered noise. Importantly, the expression of a deterministic bifurcation state that manifested subthreshold oscillations in the models enhanced their propensity to exhibit valid oscillatory traces in the presence of noise. These observations outlined the importance of interactions between different ion channels in mediating the stochastic bifurcations mediating oscillatory patterns. Furthermore, these observations implied that heterogeneities in ion-channel properties in a population of neurons would translate to heterogeneities in the emergence of bifurcation states across neurons. The strength, form, and the specific instance of noise, and its interactions with heterogeneous bifurcation states across neurons, added additional layers of variability in how these oscillatory patterns emerged. Whereas intrinsic stochasticity contributed to characteristic variability in amplitude and frequency of these oscillatory patterns, ion-channel heterogeneities translated to the pronounced neuron-to-neuron variability in all aspects of peri-threshold oscillatory patterns. The expression of peri-threshold oscillations under *in vitro* conditions could be because of ion-channel noise driving the stochastic bifurcations (White et al., 1998; Dorval and White, 2005; Fernandez and White, 2008; Fernandez et al., 2015). On the other hand, as noise levels are typically higher under *in vivo* conditions, the expression of stochastic resonance is consistent with the absence of peri-threshold oscillations under *in vivo* conditions (Fernandez and White, 2008; Schmidt-Hieber and Hausser, 2013).

Together, using a combination of theoretical, computational, and electrophysiological methods, we argue for heterogeneous stochastic bifurcations as a unifying framework that fully explains peri-threshold activity patterns in MEC stellate cells. Within this framework, we argue that the manifestation of intrinsic oscillatory patterns in MEC stellate cells should be considered as state-dependent, as several factors govern their emergence. These factors include heterogeneities in ion-channel composition and intrinsic properties, the overall synaptic drive which also drives the bifurcation parameter, the specific levels of different forms of noise, neuromodulatory tone under different behavioral state, activity-dependent neural plasticity, and channelopathies.

## MATERIALS AND METHODS

### Electrophysiology: Brain slice preparation and whole-cell current clamp recordings

#### Ethical approval

All experiments reported in this study were performed in strict compliance with the protocols cleared by the Institutional Animal Ethics Committee (IAEC) of the Indian Institute of Science, Bangalore.

For all the electrophysiological experiments, male Sprague Dawley rats of age 5–9 weeks were used. The rats were maintained in 12 hours light – 12 hours dark cycle. Food and water were provided *ad libitum*. Surgical and *in vitro* electrophysiology procedures followed previously established protocols in the laboratory (Ashhad and Narayanan, 2016; Mishra and Narayanan, 2020, 2021) and are detailed below. Rats were anesthetized by intraperitoneal injection of a mixture of ketamine-xylazine. The onset of deep anesthesia was assessed by the cessation of toe-pinch response. After deep anesthesia, transcardial perfusion was performed with ice cold cutting solution (composition: 2.5 mM KCl, 1.25 mM NaH_2_PO_4_, 25 mM NaHCO_3_, 0.5 mM CaCl_2_, 7 mM MgCl_2_, 7 mM dextrose, 3 mM sodium pyruvate, and 200 mM sucrose; pH 7.3; Osmolarity: ~300 mOsm; and saturated with 95% O_2_ and 5% CO_2_). Decapitation was performed in the presence of the cutting solution and the brain was removed quickly. Near-horizontal middle entorhinal cortical (bregma –6.5 mm to –5.1 mm) slices of thickness of 350 μm were obtained using a vibrating blade microtome (Leica Vibratome) while the brain was submerged in oxygenated ice-cold cutting solution. Brain slices were carefully transferred to the holding chamber solution (composition: 125 mM NaCl, 2.5 mM KCl, 1.25 mM NaH_2_PO_4_, 25 mM NaHCO_3_, 2 mM CaCl_2_, 2 mM MgCl_2_, 10 mM dextrose, and 3 mM sodium pyruvate) and incubated for 10 mins at 34 °C, followed by incubation at room temperature for at least 45 mins. The holding chamber was continuously oxygenated with 95% O_2_ and 5% CO_2_ gas mixture.

A slice was transferred to the recording chamber and was continuously perfused with oxygenated (95% O_2_ and 5% CO_2_ gas mixture) ACSF (125 mM NaCl, 3 mM KCl, 1.25 mM NaH_2_PO_4_, 25 mM NaHCO_3_, 2 mM CaCl_2_, 1 mM MgCl_2_, and 10 mM dextrose; pH 7.3; Osmolarity: ~300 mOsm) at flow rate of ~4 mL/min. Recordings were performed under whole-cell current clamp configuration at physiological temperature of 33–35 °C, which was ensured by a inline heater that was part of a closed loop temperature control system (Harvard Apparatus). The slice was visualized using a 10× objective for locating LII MEC and then visualized under 63× water immersion objective through a Dodt contrast microscope (Carl Zeiss Axioexaminer) to visually identify a stellate cell in layer II MEC. Whole cell current clamp recordings were performed from stellate cells using an BVC-700A (Dagan) amplifier. Borosilicate glass electrodes were pulled (P-97 Flaming/Brown micropipette puller; Sutter) from capillaries (1.5 mm outer diameter and 0.86 mm inner diameter; Sutter) to yield 3–7 MΩ tip resistance. These electrodes were filled with internal solution (120 mM K-gluconate, 20 mM KCl, 10 mM HEPES, 4 mM NaCl, 4 mM Mg-ATP, 0.3 mM Na-GTP, and 7 mM K_2_-phosphocreatine; pH 7.3 adjusted with KOH; osmolarity ~300 mOsm). For experiments requiring blockade of synaptic events, a combination of synaptic receptor blockers was added in the bath solution: 10 μM 6-cyano-7-nitroquinoxaline-2,3-dione (CNQX), an AMPA receptor blocker; 10 μM (+) bicuculline and 10 μM picrotoxin, both GABA_A_ receptor blockers; 50 μM D,L-2-amino-5-phosphonovaleric acid (D,L-APV), an NMDA receptor blocker, and 2 μM CGP55845, a GABA_B_ blocker. All synaptic blockers were procured from Allied Scientific.

For identification of stellate cells, 3 properties of these cell types were considered: the distance (100–150 μm) from the edge of the slice, the size of stellate cells are larger compared to interneurons or LIII EC pyramidal neurons, and range of characteristic electrophysiological measurements (low input resistance and strong frequency selectivity in the theta-frequency range) which were computed online during the course of the experiment.

All data acquisition was done using custom written software in the Igor Pro (Wavemetrics) environment. Unless mentioned otherwise, the sampling rate was set at 40 kHz. Series resistance was continuously monitored and compensated using the bridge balance circuit in the amplifier. Cells were discarded from final analyses if the initial RMP was more depolarized than –50 mV or if the series resistance increased above 25 MΩ or if there were temperature fluctuations observed during the experiment. All the experiments were performed at the initial resting membrane potential of the specific cell, unless otherwise specified. Voltages have not been corrected for the liquid junction potential, which was experimentally measured to be ~8 mV. Analyses and plotting of electrophysiological data were performed using custom-written programs in IGOR Pro (WaveMetrics) and in MATLAB 2018a (Mathworks). Statistical analyses of electrophysiological data were executed using the R computing and statistical package (R core Team 2013).

### Electrophysiology: Subthreshold Measurements

Resting membrane potential (*V*_*RMP*_) was computed by averaging 11 trials of the membrane potential without any current injection for 100 ms. Input resistance (*R*_in_) was calculated from the steady state voltage response (after 700 ms of current injection) of the cell to the subthreshold current pulses of amplitudes spanning −50 pA to 50 pA in steps of 10 pA (Fig. 1*A*). The slope of a linear fit to this steady state voltage-current plot was assigned as the input resistance of the cell (Fig. 1*A*). To compute temporal summation ratio (*S*_α_), five alpha-excitatory postsynaptic potentials (α-EPSPs) with 50 ms interval were evoked by current injection with following dynamics:

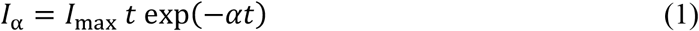

with α=0.1 ms^−1^. The temporal summation ratio (*S*_α_) was calculated as the ratio of the amplitudes of the last and first α-EPSPs (Fig. 1*B*). For estimating the sag ratio, the voltage response to a hyperpolarizing step current of 100 pA amplitude for 500 ms was recorded. Sag ratio was computed as the 100 × (1 – *V*_ss_/*V*_peak_), where *V*_ss_ and *V*_p*eak*_ denoted the steady state and peak voltage deflections from *V*_RMP_, respectively (Fig. 1*C*).

**Figure 1:**
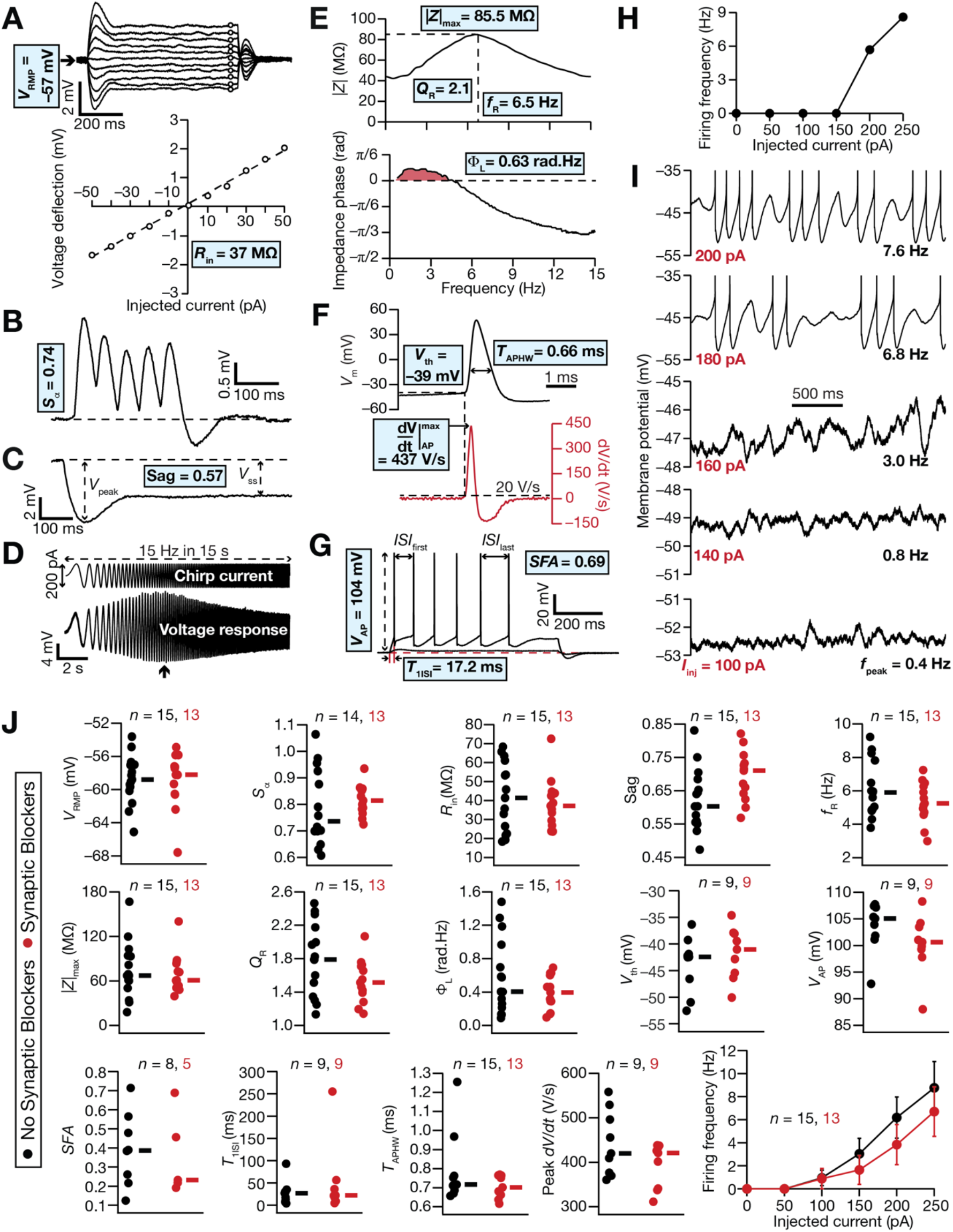
Heterogeneities in characteristic subthreshold and suprathreshold measurements from rat MEC layer II stellate cells recorded using whole-cell patch clamp electrophysiology. (A) Resting membrane potential (*V*_RMP_) was computed by averaging 11 trials of the membrane potential without any current injection with each trial spanning 100 ms. Input resistance (*R*_in_) was calculated from the steady state voltage deflection (after 700 ms of current injection) of the cell to the subthreshold current pulses of amplitudes spanning −50 pA to 50 pA in steps of 10 pA. The slope of a linear fit to this steady state voltage-current plot was assigned as the input resistance of the cell. (B) The temporal summation ratio (*S*_α_) as computed as the ratio of the last and first evoked EPSPs by five α-EPSPs separate by 50 ms interval. (C) Sag ratio (*Sag*) was computed from the voltage response to a 500 ms hyperpolarizing pulse of 100 pA amplitude, as the 100 (1 – *V*_ss_/*V*_peak_), where *V*_ss_ and *V*_peak_ denoted the steady state and peak voltage deflection from *V*_RMP_, respectively. (D) Impedance profile of neurons were computed using Chirp15 stimulus (top) which is a sinusoidal current stimulus with constant subthreshold amplitude (200 pA peak to peak) with frequency linearly spanning from 0 to 15 Hz in 15 s. The bottom panel depicts the voltage response of a neuron to the Chirp15 stimulus. The arrow depicts the temporal location where the maximal amplitude response was recorded. (E) *Top,* Impedance amplitude |*Z*| plotted as a function of frequency shown to manifest resonance. The frequency at which |*Z*(*f*)| reached its maximum value was considered as the resonance frequency, *f*_*R*_ and resonance strength (*Q*_*R*_) was defined as the ratio of |*Z*(*f*_R_)|to |*Z*(0.5)|. *Bottom,* Impedance phase plotted as a function of frequency shown to manifest positive phase values representing phase lead. The total inductive area, Φ_L_, was defined as the area under the inductive part of the impedance phase profile. (F) Action potential (AP) half-width (*T*_APHW_) was calculated as the temporal width measured at the half-maximal points of the AP peak with reference to *V*_RMP_. The maximum slope of the action potential, 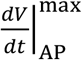, was calculated as the maximum value of the temporal derivative of the AP trace. The voltage in the AP trace corresponding to the time point at which the *dV*/*dt* crossed 20 V/s defined AP threshold *V*_th_. (G) AP amplitude (*V*_AP_) was computed as the difference between the peak voltage of the spike 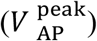 and *V*_RMP_. The ratio of duration between the first and the second spikes and that of the last and second last spike was defined as a measure of spike frequency adaption (*SFA*). (H) AP firing frequency was computed by extrapolating the number of spikes obtained during a 700 ms current injection to 1 s. The amplitude of these pulse-current injections was varied from 0 pA to 250 pA in steps of 50 pA, to construct the firing frequency *vs.* injected current (*f* – *I*) plot. (I) Peri-threshold membrane potential activity were assessed in the voltage response of the cell to depolarizing pulse current injections spanning 100–300 pA in steps of 20 pA, each lasting for 5 s. The last 3 s period of these voltage traces are depicted for 5 different current injection (*I*_inj_) values. (J) 15 characteristic electrophysiological measurements of MEC layer II stellate cells with (red) or without (black) synaptic receptor blockers in the bath.

Stellate cells of layer II of MEC are within the oscillatory network. Hence, it is essential to study the excitability to these cells with time-varying inputs of different frequencies, rather than limiting excitability measurements to pulse currents (Haas and White, 2002; Giocomo et al., 2007; Haas et al., 2007; Giocomo and Hasselmo, 2008; Dodson et al., 2011; Boehlen et al., 2013). We used chirp stimulus (Fig. 1*D*), a sinusoidal current stimulus with constant subthreshold amplitude (200 pA peak to peak) with frequency linearly spanning from 0 to 15 Hz in 15 s (Narayanan and Johnston, 2007), for characterizing the impedance profile of stellate cells. Frequency-dependent impedance, *Z*(*f*), was calculated as the ratio between the Fourier transform of the voltage response to the chirp stimulus and the Fourier transform of the chirp stimulus. The magnitude of this complex quantity defined the impedance amplitude profile (Fig. 1*E*):

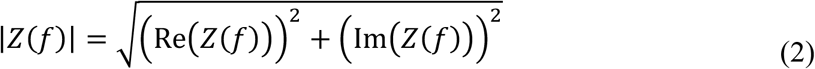

where Re(*Z*(*f*)) and Im(*Z*(*f*)) were the real and imaginary parts of *Z*(*f*), respectively. maximum impedance amplitude (|*Z*|_max_, Fig. 1*E*) was defined as the maximum value of impedance across all frequencies. The frequency at which |*Z*(*f*)| reached its maximum value was considered as the resonance frequency (*f*_R_). Resonance strength (*Q*_R_) was defined as the ratio of |*Z*(*f*_R_)| to |*Z*(0.5)| (Fig. 1*E*). The impedance phase profile was computed as (Fig. 1*E*):

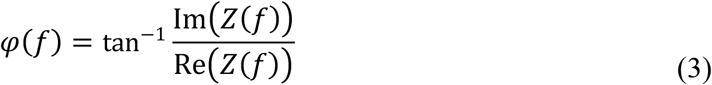

The total inductive area, Φ_L_, defined as the area under the inductive part (Fig. 1*E*) of *φ*(*f*), was calculated based on the impedance phase profile (Narayanan and Johnston, 2008):

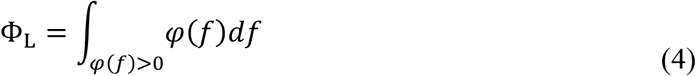

### Electrophysiology: Suprathreshold Measurements

Action potential (AP) firing frequency was computed by extrapolating the number of spikes obtained during a 700 ms current injection to 1 s. The amplitude of these step-like current injections (*I*_*inj*_) was varied from 0 pA to 250 pA in steps of 50 pA, to construct the firing frequency *vs.* injected current (*f* – *I*) plot (Fig. 1*H*). Several AP related measurements were derived from the voltage response of the cell, resting at *V*_RMP_, to a 250 pA pulse-current injection (Fig. 1*F–G*). AP half-width (*T*_APHW_) was the temporal width measured at the half-maximal points between the AP peak and *V*_RMP_ (Fig. 1*F*). The peak slope of the action potential 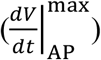 was calculated from the maximum temporal derivative of the AP trace (Fig. 1*F*). The voltage in the AP trace corresponding to the time point at which the d*V*/d*t* crossed 20 V/s defined AP threshold, *V*_th_ (Fig. 1*F*). AP amplitude (*V*_AP_) was the maximal depolarizing deflection of the AP from *V*_RMP_ (Fig. 1*G*). The temporal distance between the timing of the first spike and the onset of the pulse-current injection was defined as latency to first spike, *T*_1ISI_ (Fig. 1*G*). The ratio of duration between the first and the second spikes (the first interspike interval) and last and second last spike (the last interspike interval) was defined as a measure of spike frequency adaptation, *SFA* (Fig. 1*G*).

### Electrophysiology: Peri-threshold Activity Patterns

Stellate cells manifest intrinsic peri-threshold membrane potential oscillations with depolarizing current injections (Alonso and Llinas, 1989; Klink and Alonso, 1993, 1997; Haas and White, 2002; Giocomo et al., 2007; Haas et al., 2007; Giocomo and Hasselmo, 2008; Dodson et al., 2011; Boehlen et al., 2013). Peri-threshold membrane potential oscillations were assessed in the voltage response of the cell to depolarizing pulse current injections (Fig. 1*I*) spanning 100–300 pA in steps of 20 pA, each lasting for 5 s. Due to the intrinsic heterogeneities, every cell manifested oscillations at different peri-threshold membrane potentials. Therefore, we used this short-pulse protocol for visually identifying the range of membrane potentials where peri-threshold oscillations manifested in individual cells. The holding voltage was then maintained around the identified range for each cell and multiple 15 s long voltage responses were recorded. The long-pulse protocol provided traces that were then employed to quantitatively assess the manifestation of valid oscillatory traces.

### Stochastic Hopf Bifurcation Simulations

Neurons are often considered as high-dimensional non-linear dynamical systems (Ermentrout and Terman, 2010; Izhikevich, 2010). Upon receiving increasing amount of injected current, neurons typically switch from a stable resting membrane potential to a state where they manifest robust subthreshold oscillations or regular action potential firing. This property can be modeled by a nonlinear dynamical system that switches from a stable equilibrium point to manifesting limit cycles upon changes in a bifurcation parameter (injected current in the case of neurons). In this study, we employed a simple nonlinear dynamical system with such characteristics to assess the impact of stochasticity on robustness and spectral characteristics of the emergent oscillations. We specifically used the supercritical Poincare-Andronov-Hopf’s bifurcation (referred to as Hopf bifurcation), a non-linear dynamical system capable of showing bifurcation behavior resulting in stable limit cycles. The dynamics of Hopf bifurcation were governed by the following set of coupled differential equations:

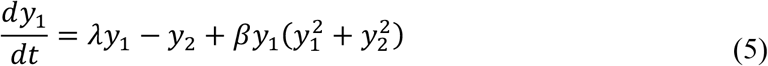

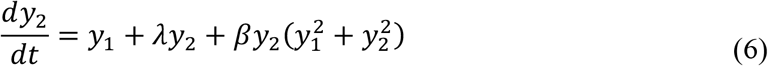

where *y*_1_ and *y*_2_ were state variables that evolved temporally in a coupled manner. The dynamics followed the supercritical Hopf bifurcation if β = −1, with equilibrium at the origin (*y*_1_ = *y*_2_ = 0). *λ* is the bifurcation parameter, with the system showing asymptotically stable dynamics for *λ* ≤ 0 and manifesting stable limit cycles for *λ* > 0. We set the integration time constant for the Hopf bifurcation simulations at 25 μs (40 kHz sampling rate) to match with the integration time constant of neuronal models.

Equations (5)–(6) define a deterministic Hopf bifurcation. To assess the impact of noise on this dynamical system, we introduced stochasticity as either *extrinsic* (external) or *intrinsic* (parametric) noise. Extrinsic noise was introduced as additive Gaussian white noise (GWN) to the equations governing the Hopf bifurcation:

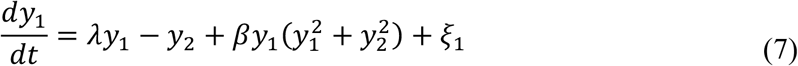

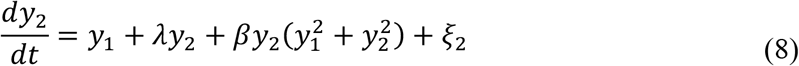

where, *ξ*_1_ and *ξ*_2_ were independent, identically distributed, and were drawn at every time step from normal distributions with variance 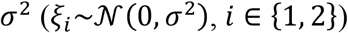. The level of noise was varied by adjusting *σ*^2^. Intrinsic stochasticity was introduced by introducing additive GWN to the bifurcation parameter *λ* that governs the emergence of stable limit cycle. Specifically, the equations governing a stochastic Hopf bifurcation with parametric noise were:

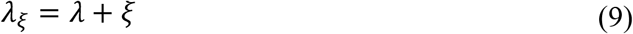

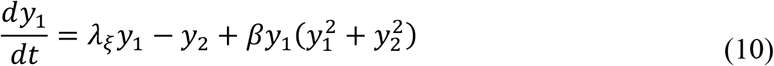

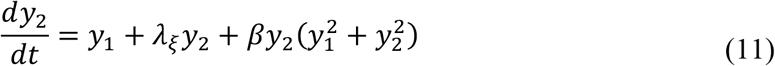

where *ξ* was drawn from a normal distribution with variance 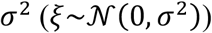. The parameter *λ*_*ξ*_ was updated at each time step of the simulation, and the level of noise was varied by adjusting *σ*^2^.

### Heterogeneous population of MEC layer II stellate cell models

We used 155 conductance-based heterogeneous population of medial entorhinal cortex (MEC) layer II stellate neuronal models constructed previously (Mittal and Narayanan, 2018). The methodology for the generation of these 155 models and the measurements employed for their validation as MEC LII stellate neurons are as detailed before (Mittal and Narayanan, 2018). Briefly, each neuron in this heterogeneous population was designed as a single compartment cylindrical model of length 75 μm and diameter 70 μm. The passive properties of the model were defined by a specific membrane resistance (*R*_m_) that determined the leak-channel density and a specific membrane capacitance (*C*_m_). A set of nine distinct voltage- or calcium-gated ion channel subtypes, whose gating properties and kinetics were derived from MEC LII stellate cells, controlled the active properties of these neuronal models (Mittal and Narayanan, 2018). These channels were fast sodium (NaF), delayed rectifier potassium (KDR), hyperpolarization-activated cyclic-nucleotide gated (HCN) nonspecific cationic, persistent sodium (NaP), *A*-type potassium (KA), low-voltage-activated (LVA) calcium, high-voltage-activated (HVA) calcium, *M*-type potassium (KM), and small-conductance calcium-activated potassium (SK) channels. All channel kinetics were based on the Hodgkin-Huxley formulation (Hodgkin and Huxley, 1952) for ion channel models, except for the SK channel, which followed six-state Markovian model (Mittal and Narayanan, 2018). The increase in cytosolic calcium was defined by voltage-gated calcium channel activity and decayed following a first order kinetics with a decay time constant of τ_Ca_.

We had employed a multi-parametric multi-objective stochastic search (MPMOSS) algorithm for identifying this heterogeneous population of 155 biophysically valid MEC layer II stellate cell model (Mittal and Narayanan, 2018). The stochastic search was performed over 55 parameters, which governed the passive and active properties of the neurons. The validation process required that the model satisfied multiple objectives, specifically requiring 10 characteristic electrophysiological measurements to fall within their respective experimental ranges. The 10 subthreshold and suprathreshold *in vitro* measurements from MEC LII stellate cells were resting membrane potential (*V*_R*MP*_), standard deviation of the resting membrane potential (*V*_SD_), sag ratio (*Sag*), input resistance (*R*_in_), resonance frequency (*f*_R_), resonance strength (*Q*_R_), subthreshold oscillation frequency (*f*_osc_), number of action potential elicited for a 100 pA (*N*_100_) or 400 pA (*N*_400_) current injection, and amplitude of action potential (*V*_AP_). In performing the stochastic search process, we randomly picked a value for each of 55 parameters from their respective uniform distribution to construct a single model neuron. This model was considered a “valid” neuronal model if it passed all the multiple objectives involving the 10 electrophysiological measurements and was rejected otherwise. This procedure of random selection followed by validation of models was repeated 50,000 times and had yielded a heterogenous population of 155 valid models (Set I in (Mittal and Narayanan, 2018)). These 155 models manifested theta-frequency subthreshold membrane potential oscillations in the theta-frequency range (3 ≤ *f*_osc_ ≤ 10 Hz) as they were validated with *f*_osc_ as one of the criteria (Mittal and Narayanan, 2018). These 155 valid models that satisfied all the 10 validation criteria were referred to as θ+ models (Fig. 8*A*).

Of the 50,000 models that were tested, there were a total of 623 models (including the 155 θ+ models) which satisfied 9 of the 10 characteristic electrophysiological measurements, sparing the 10^th^ criterion on intrinsic subthreshold oscillations (*f*_osc_). *f*_osc_ was computed as the frequency at the maximum power of the Fourier spectrum of the entire trace, for a subthreshold voltage trace near action potential threshold (Mittal and Narayanan, 2018). Of these 623 deterministic models, 468 models did not manifest theta-frequency subthreshold membrane potential oscillations. We randomly picked 155 models from this set of 468 models — which satisfied 9 characteristic electrophysiological properties of but did not manifest subthreshold intrinsic oscillations — for further analysis and called them as θ– models (Fig. 8*A*).

### Introducing stochasticity into the heterogeneous population of MEC layer II stellate cell models

The heterogeneous MEC LII stellate cell models were deterministic in nature, with the dynamics entirely driven by the fixed active and passive channel properties of the specific neuronal model. However, biological neurons are inherently stochastic, with two predominant forms of noise governing the function of individual neurons (Faisal et al., 2008): (i) ion-channel noise, resultant from the stochastic nature of the gating properties of individual ion channels; and (ii) synaptic noise, consequent to background synaptic activity and the stochastic nature of synaptic function. We introduced ion-channel noise by making the activation gating parameter (*m*_N*ap*_) of NaP channel in the model to be stochastic. Specifically, a zero mean GWN was added to the value of *m*_N*ap*_ at every time step and the level of noise was governed by the value of standard deviation of the GWN. Synaptic noise was introduced through a balanced high conductance state, which was introduced such that the average resting membrane potential was not altered in the presence of balanced excitatory and inhibitory synapses impinging on the model neuron. The balance was achieved by adjusting the number and frequency of activation of the different excitatory (*N*_e_ = 100, each with an average firing frequency of 3 Hz) and inhibitory (*N*_i_ = 20, each with an average firing frequency of 10 Hz) synapses. Both excitatory and inhibitory synaptic conductances were modelled as the sum of two exponentials, one defining the rise (τ_r_= 2 ms) and other governing the decay (*τ*_*d*_= 10 ms). The reversal potential for excitatory and inhibitory synapses were 0 mV and –80 mV, respectively. Different levels of synaptic noise were achieved by increasing the value of scalar multipliers to the synaptic conductance values. A third form of stochasticity was introduced as extrinsic additive noise was introduced by injection of a zero mean GWN current into the model, with noise level adjusted by altering the standard deviation.

### Assessment of peri-threshold oscillations in the stochastic heterogeneous population of models

The protocol for generating peri-threshold activity traces of a given model, for assessment for the expression of membrane potential oscillations, involved step current injections ranging from 100 pA to 300 pA in steps of 10 pA, for 5 s each (21 traces generated by the protocol). This protocol was repeated for all noise levels (5 levels for each of additive and ion-channel noise and 4 levels for synaptic noise), for all 3 sources of noise, across each of the 155 θ+ and the 155 θ– models (21×(5+5+4)×(155+155)=91,140 traces). This entire process was repeated for 10 unique trials with different seed values for the generation of noise, yielding a total of 911,400 traces each of 5 s duration. In addition, there were 21 traces for each level of current injection from the deterministic (no noise) version of these 155 θ+ and the 155 θ– models, yielding a further 6510 traces. Together, these yielded a total of a requirement of validating 917,910 model traces.

We used these 5 s voltage recordings for quantitative analyses of peri-threshold oscillatory activity. While using Fourier transform or Lomb’s periodogram, we let the simulations to settle by removing the initial 2 s of neuron response to current injection and performed all the analyses for last 3 s of voltage traces. For comparison purposes, spectral analyses of traces from Hopf bifurcation simulations were also performed over the last 3 s of the time series. We used wavelet transform to perform time-frequency analysis of outputs from the Hopf bifurcation simulations, from the neuronal models, and from electrophysiological recordings.

Each of the 911,400 traces from the stochastic versions of the θ+ and θ– models were individually assessed for the expression of robust oscillations. Traces were declared as valid oscillatory traces if they satisfied all five measurements derived from their wavelet spectra. Two measurements were derived from the frequency values at maximal power computed from the wavelet spectrogram. Specifically, at each time point, we noted down the frequency at which maximal power was observed, yielding an array of frequency values of the same duration as the original trace (3 s). The mean frequency, μ_fmax_ of these set of frequency values was required to be within the theta frequency range (3–10 Hz). We used the standard deviation, σ_fmax_ of these set of frequency values as a measure of frequency variability within the trace and required this to be < 1 Hz for frequency stability.

Two other measurements placed constraints on the power at the mean frequency. Here, we noted down the power at the mean frequency (μ_fmax_ computed earlier) at each time point, yielding an array of power values of the same duration as the original trace (3 s). The mean power, μ_Pfmean_ computed from this array was required to be > 0.5 (a.u.), setting a threshold on the minimal power at the mean frequency of oscillations. The standard deviation, σ_Pfmean_ computed from this array was a measure of how variable the power at the mean frequency was within the trace and was required to be <1 (a.u.) as a measure of power stability. The fifth measurement that we defined to validate oscillatory traces was spirality coefficient (ζ), computed as the slope of a linear fit of the plot of oscillatory power at the mean frequency (the array used for the power measurements) *vs*. time. The absolute value of the spirality coefficient was required to be <0.5 for valid oscillatory traces, specifically to avoid decaying or expanding oscillations.

Quantitatively, the five criteria for identifying robust theta-frequency oscillations in these traces were: 3 < μ_fmax_ < 10 Hz; σ_fmax_ < 1 Hz, μ_Pfmean_ > 0.5; σ_Pfmean_ < 1; |ζ| < 0.5. These criteria and bounds on each of them were used to validate oscillations in traces from stochastic Hopf bifurcations and from theta-filtered noise, as well as activity patterns from stochastic MEC stellate cell models and electrophysiological recordings from rat MEC stellate cells.

### Computational details

All simulations involving conductance-based stellate cell models were performed using the NEURON programming environment (Carnevale and Hines, 2006) at 34 °C, with a simulation step size of 25 μs. All Hopf bifurcation simulations and generation of theta-filtered noise traces were performed in MATLAB 2018a (Mathworks). All data analyses and plotting were implemented using custom-written scripts on MATLAB 2018a (Mathworks) or IGOR Pro (WaveMetrics) environments. All statistical analyses were performed using the R computing and statistical package (R core Team 2013).

## RESULTS

The central hypothesis assessed in this study is that peri-threshold activity patterns observed in medial entorhinal cortex (MEC) LII stellate cells are consistent with those elicited by stochastic bifurcations in a heterogeneous neuronal population. An ongoing debate in the field considers these activity patterns either as rhythmic oscillations or as theta-filtered noise. We argue that the activity patterns observed in MEC stellate cells are consistent with neither. Instead, we provide several lines of evidence for these activity patterns being consistent with the dynamics of a heterogeneous population of nonlinear systems manifesting stochastic bifurcations.

### Electrophysiological recordings from rat MEC stellate cells manifested intrinsic peri-threshold oscillations and unveiled heterogeneities in characteristic physiological properties

We performed two sets of whole-cell current-clamp intracellular recordings from visually identified rat MEC stellate cells, in the presence or absence of synaptic receptor blockers. We characterized MEC stellate cells using several measurements (Fig. 1*A–I*) and confirmed that they manifested signature electrophysiological properties (Fig. 1*J*): depolarized resting membrane potential (*V*_R*MP*_), large sag potential (*Sag*), low excitability (*R*_in_, |*Z*|_max_, and firing frequency) and temporal summation (*S*_α_), strong frequency selectivity in the theta-frequency range (3 < *f*_R_ < 10 Hz; high *Q*_R_), and importantly, the manifestation of peri-threshold oscillations (Fig. 1*I*, Figs. S1–S2). We found that the intrinsic properties of MEC stellate cells manifested pronounced heterogeneities, irrespective of whether recordings were performed in the presence (*n* = 13) or (*n* = 15) absence of synaptic blockers in the bath (Fig. 1*J*; Tables S1–S2). These electrophysiological recordings confirmed earlier computational predictions (Mittal and Narayanan, 2018) that the impedance phase should manifest a lead in the lower frequency ranges, with the total inductive phase (Φ_L_) showing nonzero values (Fig. 1*E*; Fig. 1*J*).

These intrinsic measurements were dependent on the voltage at which recordings were performed (Fig. S3*A*) and showed weak correlations across most pairs of measurements from all stellate cells (*n* = 28) that were recorded from (Fig. S3*B–C*). Some of the measurements that showed relatively strong correlations are known to be dependent on the same sets of ion channels (Mittal and Narayanan, 2018). These weak correlations across most of the pairs confirmed that the constellation of measurements employed to characterize MEC stellate cells were providing insights about different aspects of stellate cell physiology.

As our focus here was on peri-threshold oscillations, we used two distinct set of electrophysiological protocols to record peri-threshold activity patterns. First, we used a short-pulse protocol to identify the peri-threshold voltage range for each neuron (Fig. 1*I*, Fig. S1). The holding voltage was then maintained around the identified range for each cell and multiple 15 s long voltage responses were recorded (Fig. S2). Qualitatively, we observed peri-threshold oscillations with both short as well as long protocols (Fig. 1*I*, Fig. S1–S2). Importantly, we also found instances of spike clustering, a signature characteristic of MEC stellate cells (Alonso and Klink, 1993; Dickson et al., 2000b; Fransen et al., 2004; Pastoll et al., 2012), in peri-threshold mixed-mode oscillatory traces (Fig. 1*I*, Fig. S1–S2).

Together, electrophysiological recordings from rat MEC stellate cells unveiled pronounced cell-to-cell variability in characteristic physiological measurements (Fig. 1, Fig. S3), and showed the manifestation of characteristic subthreshold oscillations or mixed-mode oscillations with spike clustering in the peri-threshold voltage ranges.

### The dynamics of a nonlinear system manifesting stochastic bifurcations were consistent with peri-threshold activity patterns of MEC stellate cells

An ongoing debate on the peri-threshold intrinsic activity patterns manifested by MEC LII stellate cells (Fig. 1*I*, Fig. S1–S2) argue for them to be rhythmic oscillatory patterns akin to a sinusoid (Fig. 2*A*) or consider them to be filtered noise (Fig. 2*B*). Different signal processing tools, including Fourier analysis, Lomb’s periodogram analysis, and wavelet spectrogram (Fig. 2*A–B*), have been employed to assess these activity patterns for their consistency with either of these frameworks (Dickson et al., 2000a; Dickson et al., 2000b; Erchova et al., 2004; Giocomo et al., 2007; Nolan et al., 2007; Giocomo and Hasselmo, 2008; Dudman and Nolan, 2009; Giocomo and Hasselmo, 2009; Dodson et al., 2011; Yoshida et al., 2011; Pastoll et al., 2012; Boehlen et al., 2013). The manifestation of periodic oscillations has important implications for several network models involving these neurons in the emergence of grid cell firing. Motivated by the specific equivalence of neurons in general (Ermentrout and Terman, 2010; Izhikevich, 2010), and LII stellate cells in particular (White et al., 1995; White et al., 1998; Dickson et al., 2000a; Dickson et al., 2000b; Acker et al., 2003; Erchova et al., 2004; Fernandez et al., 2015), to nonlinear systems manifesting bifurcations yielding stable limit cycles, here we argue that their activity patterns are consistent with systems manifesting stochastic bifurcations. To systematically build our argument, we picked the Poincare-Andronov-Hopf bifurcation (equations 5–6), a simple nonlinear dynamical system capable of manifesting a deterministic bifurcation yielding stable limit cycles. This deterministic nonlinear system manifests an equilibrium point at the origin and switches between a no-oscillation state involving a stable spiral to an oscillatory state manifesting stable limit cycles with changes in a bifurcation parameter.

**Figure 2:**
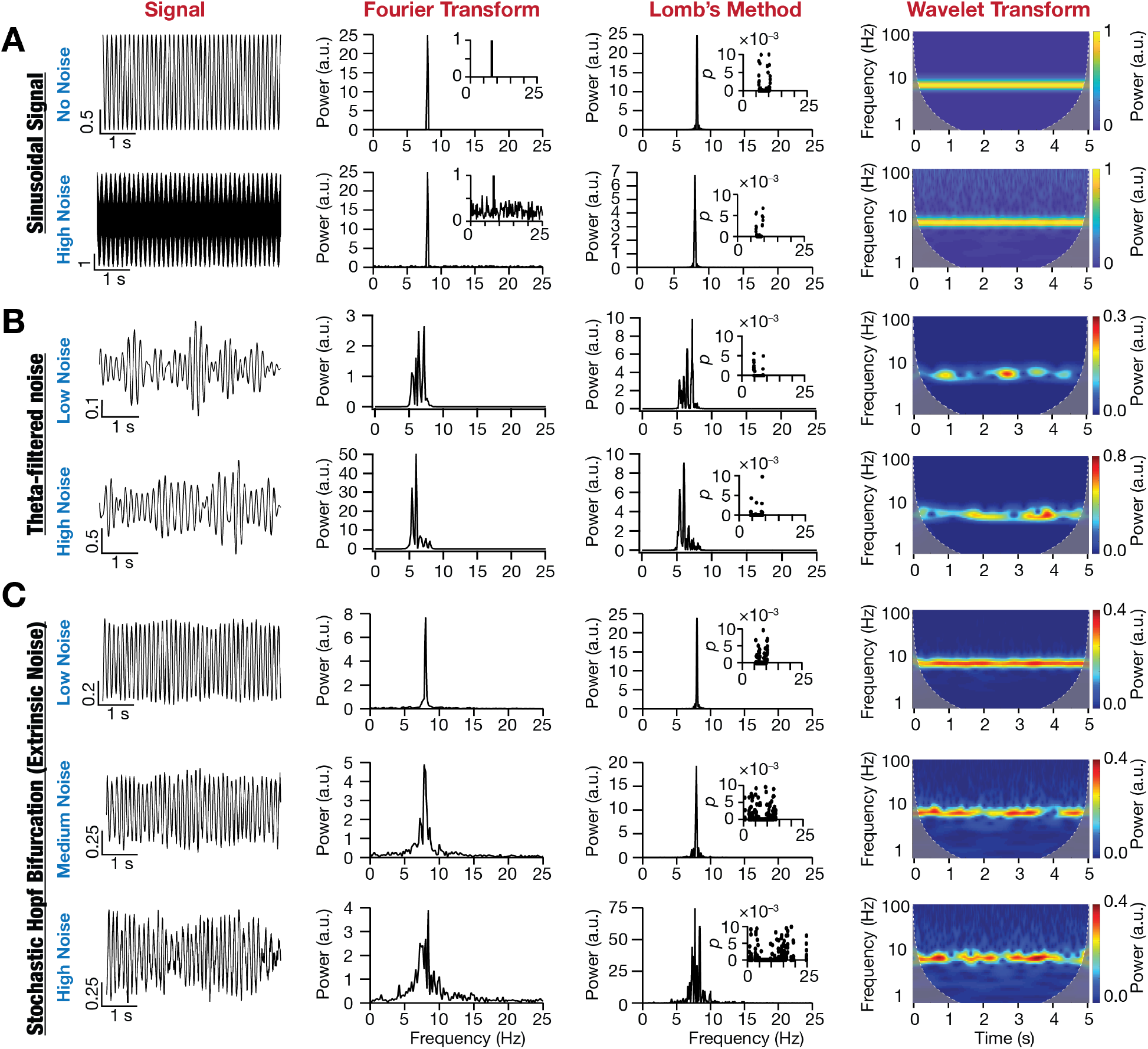
Spectral properties of a noisy sinusoidal signal, of filtered noise signals, and of oscillations emergent from a stochastic non-linear dynamical system. (A) Impact of zero mean Gaussian white noise (GWN) on the frequency content of an 8 Hz sinusoidal signal. *Top row*: No noise, *Bottom row*: High noise. *Column 1*: time-domain signal, *Column 2*: Fourier Transform (notice the 8 Hz peak) of the signal, *Column 3*: Lomb’s periodogram of the signal with inset depicting the significance of each peak in the periodogram, and *Column 4*: spectrogram of the signal computed using wavelet transform. (B) Impact of altering the variance on filtered zero mean Gaussian white noise traces. *Row 1*: Low noise and *Row 2*: High noise. *Columns 1–4*: same as panel A. (C) Impact of additive zero mean GWN (extrinsic noise) on oscillations emerging from a nonlinear dynamical system (Hopf bifurcation). *Row 1*: Low noise, *Row 2*: Medium noise, and *Row 3*: High noise. *Columns 1–4*: same as panel A.

We introduced stochasticity by means of extrinsic (equations 7–8) or intrinsic (equations 9–11) perturbations, thereby introducing the ability of the system to stochastically switch between different bifurcation states, thus exhibiting stochastic bifurcations. We demonstrated that the dynamics of this simple system manifesting stochastic bifurcations, with either extrinsic (Fig. 2*C*) or intrinsic (Fig. S4) noise, were qualitatively and quantitatively consistent with activity patterns from MEC LII stellate cells. Specifically, qualitatively, we noted that these traces exhibited oscillations that were irregular, non-rhythmic, were of variable amplitude, and noisy (Fig. 2*C*; Fig. S4). Quantitatively, we observed multiple peaks in their Fourier spectra, multiple statistically significant peaks in their Lomb’s periodogram, and their wavelet spectrograms reflected the variable amplitude structure (Fig. 2*C*, Fig. S4). In the absence of noise, the deterministic Hopf bifurcation generated rhythmic oscillatory patterns, translating to a single sharp peak in the spectral domain. However, the introduction of stochasticity, either in the bifurcation parameter or as an external perturbation, the stochastic Hopf bifurcation elicited variable amplitude and arhythmic patterns of activity that were qualitatively and quantitatively consistent with subthreshold oscillations of LII stellate cells (Fig. 1*I*, Fig. S1–S2).

### Different forms of noise stabilized peri-threshold oscillatory activity in MEC stellate cell models

Motivated by the ability of a simple system manifesting stochastic bifurcations to show activity patterns that were consistent with MEC stellate cell activity, we next turned to introducing stochasticity into conductance-based models of MEC stellate cells. Stellate cells of the MEC could be considered as a multi-dimensional non-linear dynamical system endowed with multiple bifurcation states achieved with increasing values of injected current: resting state, subthreshold oscillations, mixed-mode oscillations (involving subthreshold oscillations and action potential firing), regular action potential firing, and depolarization-induced block. Specific sets of ion channels and their expression profiles, characteristic passive properties, and interactions among these different components together allow these neurons to manifest these signature bifurcation states (Alonso and Llinas, 1989; Alonso and Klink, 1993; Klink and Alonso, 1993; White et al., 1993; Dickson et al., 2000b; Fransen et al., 2004; Dorval and White, 2005; Dodson et al., 2011).

As MEC stellate cells manifest pronounced heterogeneities in their parameters and intrinsic properties (Fig. 1), including cell-to-cell variability in the precise values of the injected current required for these bifurcation states (Alonso and Llinas, 1989; Alonso and Klink, 1993; Klink and Alonso, 1993; White et al., 1993; Dickson et al., 2000b; Fransen et al., 2004; Dorval and White, 2005; Dodson et al., 2011), it was essential that we employed a heterogeneous population of MEC stellate cell models. The use of a single hand-tuned model would have yielded a model with a specific form of oscillatory pattern and would not have been endowed with the biophysical heterogeneities that are intrinsic MEC stellate cells. Interpretations from a single hand-tuned model would thus be biased by the specific choice of parameters that were introduced in that single model. To avoid these biases, we employed a heterogeneous population of 155 models that was generated previously (Mittal and Narayanan, 2018) through an unbiased search and was validated against 10 different characteristic physiological properties of these MEC stellate cells. Importantly, the ranges of measurements and the interdependence among measurements obtained by electrophysiologically recorded stellate cells (Fig. 1, Fig. S3) were comparable to their counterparts in this heterogeneous stellate cell model population (Mittal and Narayanan, 2018). These 155 models also exhibited heterogeneities in each of the 55 parameters that governed ion-channel, passive, and calcium-handling properties of the model (Mittal and Narayanan, 2018).

Although these deterministic models manifested membrane potential oscillations in the theta-frequency range at certain peri-threshold voltage ranges, there were certain models that exhibited unphysiological activity patterns at specific current injection values. For instance, there were deterministic models that manifested unphysiological inward (Fig. 3*A*) or outward (Fig. 3*C*, 3*D*) spirals, yielding unstable membrane potential oscillations. We introduced three different forms of stochasticity into this heterogeneous population of MEC stellate cell models to assess the ability of noise to stabilize peri-threshold oscillations in these models. Ion-channel noise was introduced to mimic intrinsic stochasticity in ion-channel gating properties. Synaptic noise impinged on the neuronal model as fluctuations imposed by balanced excitatory and inhibitory conductance-based synapses. Additive noise was injected as a current into the model to mimic external perturbations.

**Figure 3:**
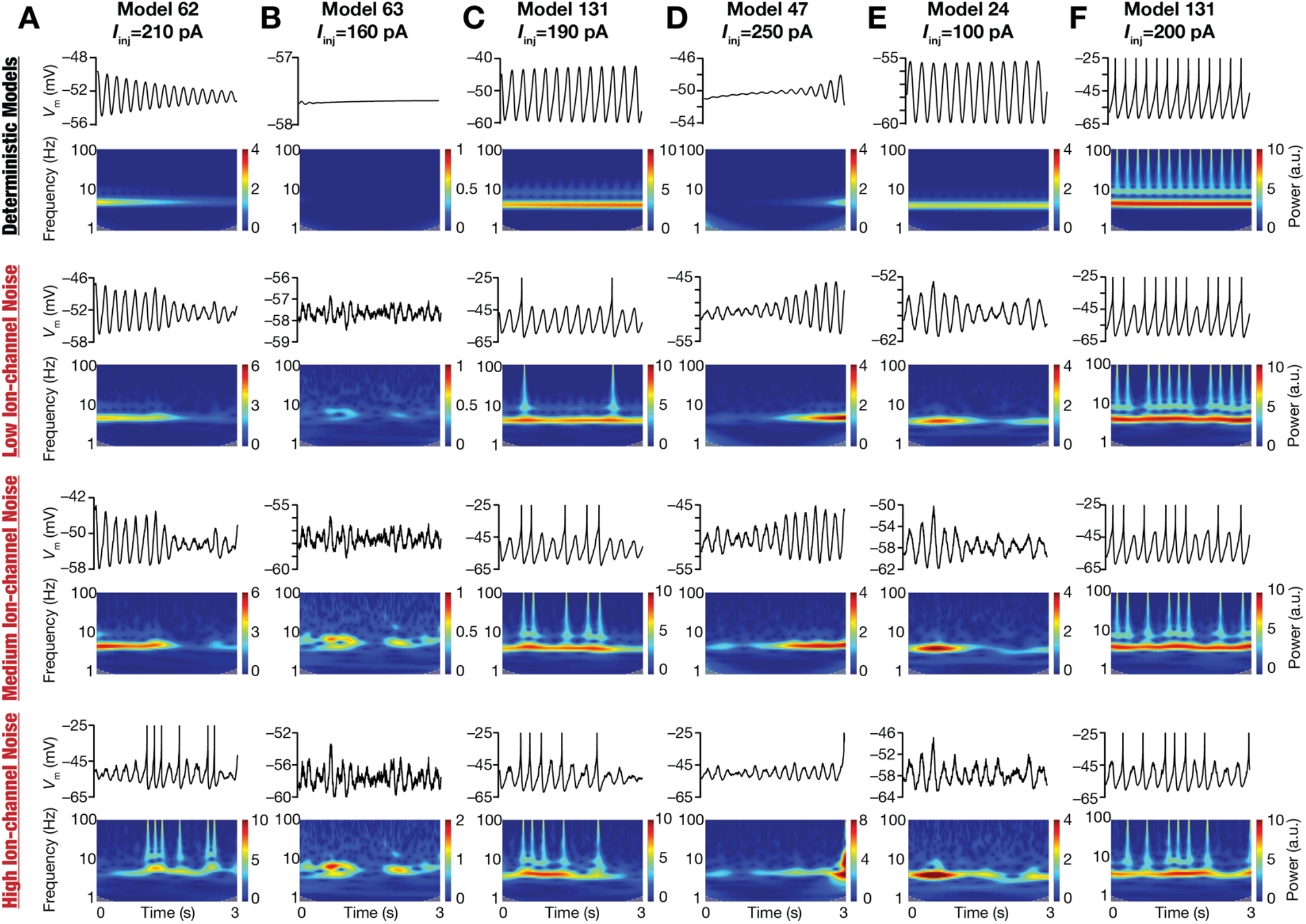
Illustrative examples of the role of ion-channel noise in stabilizing peri-threshold oscillatory patterns in a heterogeneous population of LII MEC stellate cell models. Each row in panels (A–F) depicts intrinsic activity patterns from different models for a 3 s period and the corresponding spectrograms computed using wavelet transform. Each column is identified by the corresponding model number along with identified values of injected current (*I*_inj_) employed to generate the activity patterns. The first row depicts activity patterns in the deterministic model, in the absence of any form of noise. Rows 2–4 depict activity patterns from the same model, injected with the same *I*_inj_ value, with low (0.3), medium (1.2), and high (4.8) ion-channel noise. Note when spikes occurred, they were truncated to −25 mV to emphasize the subthreshold dynamics.

The impact of these three forms of noise on a specific set of models manifesting different kinds of peri-threshold activity patterns provided important insights about the stabilizing role of stochasticity (Fig. 3, Figs. S5–S6). Specifically, deterministic models that manifested unphysiological and unstable oscillations that were decaying (Fig. 3*A*, Fig. S5*A*, Fig. S6*A*) or expanding (Fig. 3*C*–*D*, Fig. S5*C*–*D*, Fig. S6*C*–*D*) in amplitude switched to showing non-rhythmic, variable amplitude oscillations that were similar to those observed in MEC stellate cells. Deterministic models that did not manifest oscillatory patterns for a specific current injection manifested non-rhythmic, variable amplitude oscillations with the introduction of noise (Fig. 3*B*, Fig. S5*B*, Fig. S6*B*). Importantly, deterministic models that manifested regular subthreshold oscillations also switched to showing non-rhythmic, variable amplitude oscillations (Fig. 3*E*, Fig. S5*E*, Fig. S6*E*). Certain deterministic models showing purely subthreshold dynamics switched to mixed-mode oscillations at certain levels of noise (Fig. 3*A*–*D*, Fig. S5*A*–*D*, Fig. S6*A*–*D*), whereby there were spikes riding on top of subthreshold variable amplitude oscillations (Fig. 3, Figs. S5– S6). Deterministic models manifesting spiking also switched to mixed-mode oscillations with the introduction of stochasticity (Fig. 3*F*, Fig. S5*F*, Fig. S6*F*). Strikingly, models that exhibited mixed-mode oscillations manifested theta skipping in their spiking activity, also resulting in clustering of spikes (Fig. 3, Figs. S5–S6) that is observed in MEC stellate cell activity (Fig. 1*I*, Fig. S1–S2).

The rich heterogeneity of these activity patterns across models and how they emerge as functions of increasing noise levels, even with the same levels of noise across models, underscores the need to employ a heterogeneous population of models rather than employing a single hand-tuning model. The deterministic population of neurons manifested different kinds of activity peri-threshold patterns as a consequence of the heterogeneous ion-channel composition: regular subthreshold oscillations, regular spiking, no oscillations, decaying oscillations, or expanding oscillations. The strength and type of stochasticity added an additional layer of variability in how these oscillatory patterns emerged. Qualitatively, the heterogeneous population captured important stabilizing roles for stochasticity in yielding intrinsic oscillatory patterns in MEC stellate cell models (Fig. 3, Figs. S5–S6) that were similar to their physiological counterparts: (i) the ability to trigger oscillatory patterns even when oscillations are absent in the deterministic model; (ii) the ability to convert decaying or expanding oscillations in the deterministic model to variable amplitude, non-rhythmic oscillations; (iii) the ability to transform regular subthreshold oscillations in the deterministic model to non-rhythmic, variable amplitude oscillations; and (iv) the ability to generate mixed-mode oscillations with theta-skipping and clustered spikes in deterministic models showing subthreshold or suprathreshold activity patterns. The specific intrinsic activity patterns and the switches between the different bifurcation states in these model neurons were driven by their intrinsic heterogeneities, the value of the bifurcation parameter (the amount of injected current), and the form and level of noise. Together, these observations point to the manifestation of heterogeneous stochastic bifurcations across the population of MEC stellate cell models.

### Development of quantitative metrics based on the wavelet spectrogram for detecting stable oscillatory activity in peri-threshold voltage traces recorded from MEC stellate cell models

Although visual inspection of activity patterns obtained with noise qualitatively suggested a stabilizing role for noise in the emergence of peri-threshold oscillations, it is essential to develop quantitative metrics for the emergence of valid oscillations. The complexity of the validation task is enormous, considering 155 different models, each assessed at 21 different levels of depolarizing current values, with several levels (5 levels for each of additive and ion-channel noise and 4 levels for synaptic noise) of three different forms of noise at each current level of each model (21×(5+5+4)×155=45,570 traces). As noise is stochastic by definition, different instances of the same level of the same form of noise could result in very different outcomes even in the same model. Thus, there was a need to obtain multiple activity patterns for each of these different cases. We performed 10 independent trials for each model at each current injection, for each level and each form of noise (with different seed values), together yielding 455,700 stochastic activity traces. In addition, there were 21 traces for each level of current injection from the deterministic (no noise) version of these 155 models, yielding a further 3255 traces. Together, these yielded a total of a requirement of validating 458,955 model traces.

To quantify this validation process, we defined five different validation criteria for metrics from the spectrogram of these voltage traces, towards detecting stable theta-frequency oscillations. Representative valid and invalid oscillatory traces that respectively passed and failed each of these five criteria are shown in Figure 4. The first metric was μ_fmax_, the mean of the frequency values corresponding to the maximal power at each time point of the spectrogram (Fig. 4*A*) and was required to be in the theta frequency range (3–10 Hz). The standard deviation, σ_fmax_ of these set of frequency values was a measure of frequency variability (Fig. 4*B*), and the criteria was σ_fmax_ < 1 Hz for frequency stability across the trace. The third validation criteria involved μ_Pfmean_, the mean of the power values computed at μ_fmax_ at each time point (Fig. 4*C*), was required to have a certain minimal power (0.5). The standard deviation, σ_Pfmean_ computed from this set of power values was a measure of power variability (Fig. 4*D*) and was required to be <1 for power stability. The final measurement that we defined to validate oscillatory traces was spirality coefficient (ζ), was used to avoid inward or outward spirals (Fig. 4*E*) and was required to be <0.5 for valid oscillatory traces. Quantitatively, voltage traces that satisfied all these five criteria were classified as valid oscillatory traces showing robust theta-frequency peri-threshold oscillations: 3 < μ_fmax_ < 10 Hz; σ_fmax_ < 1 Hz, μ_Pfmean_ > 0.5; σ_Pfmean_ < 1; |ζ| < 0.5. Importantly, these metrics were not derived from traces that were filtered in any specific frequency range, instead assessing the entire spectrogram with specific constraints on peak frequency values (μ_fmax_), thereby avoiding artifacts that emanate from filtering (*e.g.*, Fig. 2*B*).

**Figure 4:**
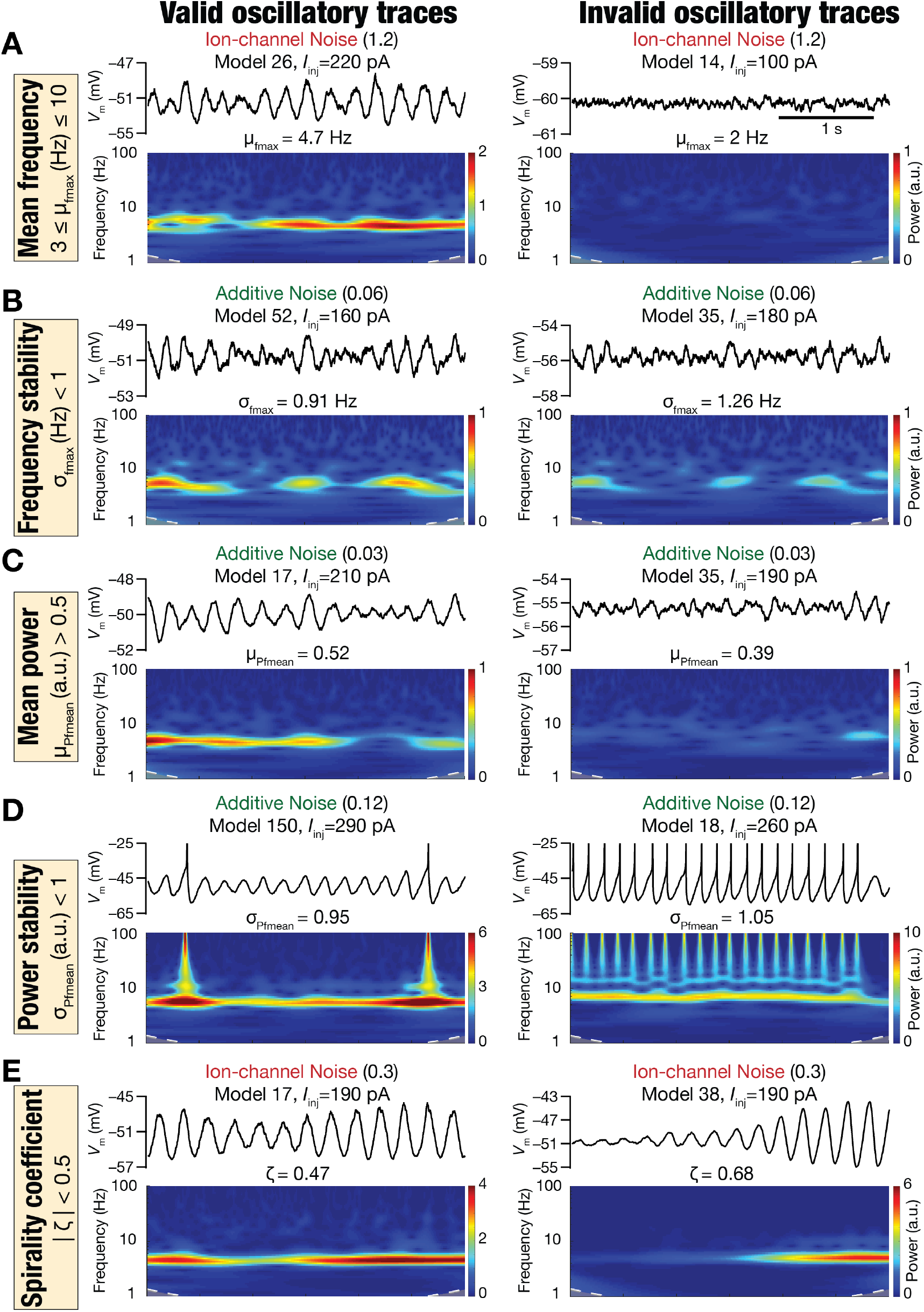
Illustration of the five spectrogram-based quantitative metrics used for assessing robustness in theta-frequency oscillatory activity with examples of valid *vs.* invalid traces from a stochastic, heterogeneous population of entorhinal stellate cells. Examples of valid (*Left*) and invalid (*Right*) oscillatory traces obtained from peri-threshold oscillatory patterns from different neuronal models at identified values of injected current (*I*_inj_). The form and the level of noise employed to generate the activity pattern are also depicted. For each case, shown are the time-domain traces (*Top*) and the respective spectrogram computed through wavelet transform (*Bottom*). The example valid and invalid traces are shown with reference to each of the five spectrogram-based quantitative metrics for assessing robustness in theta-frequency oscillatory activity: (A) Mean frequency at maximal power, μ_fmax_; (B) Standard deviation of frequency at maximal power, σ_fmax_; (C) Mean power at mean frequency, μ_Pfmean_; (D) Standard deviation of power at mean frequency, σ_Pfmean_; (E) Spirality coefficient, ζ.

### Validation of peri-threshold oscillatory traces obtained from electrophysiological recordings of rat MEC stellate cells

The validation criteria defined above were designed to eliminate non-physiological, non-theta, and weak oscillations, and imposed specific constraints on frequency and power stability. However, were these criteria sufficient to capture the variable-amplitude and irregular oscillatory patterns observed in electrophysiological recordings of MEC stellate cells? To directly assess this, we identified valid oscillatory traces from our electrophysiological recordings (Fig. 1*I*, Fig. S1–S2) using the criteria we developed based on measurements from the spectrogram: 3 < μ_fmax_ < 10 Hz; σ_fmax_ < 1 Hz, μ_Pfmean_ > 0.5; σ_Pfmean_ < 1; |ζ| < 0.5 (Fig. 5*A–B*). We detected valid peri-threshold oscillations in recordings obtained with or without synaptic blockers in all recorded MEC stellate cells (Figs. 5–6). There was pronounced neuron-to-neuron variability in the specific voltage ranges (the mean voltage of valid oscillatory traces spanned a wide range between –60 to –30 mV) where valid peri-threshold oscillations were observed (Fig. 5*C*; Fig. 6). The frequency of these oscillations also manifested heterogeneities within the theta-frequency range (3–10 Hz). The oscillations within the recorded periods were either subthreshold with the peak-to-peak amplitude <10 mV in most cases or mixed-mode oscillations that elicited action potentials in some oscillatory cycles (Fig. 5*A*, Fig. 1*I*, Fig. S1–S2). The characteristic features of these valid oscillatory traces across all recorded cells were assessed in raw traces or in traces that were median filtered to remove spikes, and manifested comparable heterogeneities in recordings obtained without or with synaptic blockers (Fig. 6).

**Figure 5:**
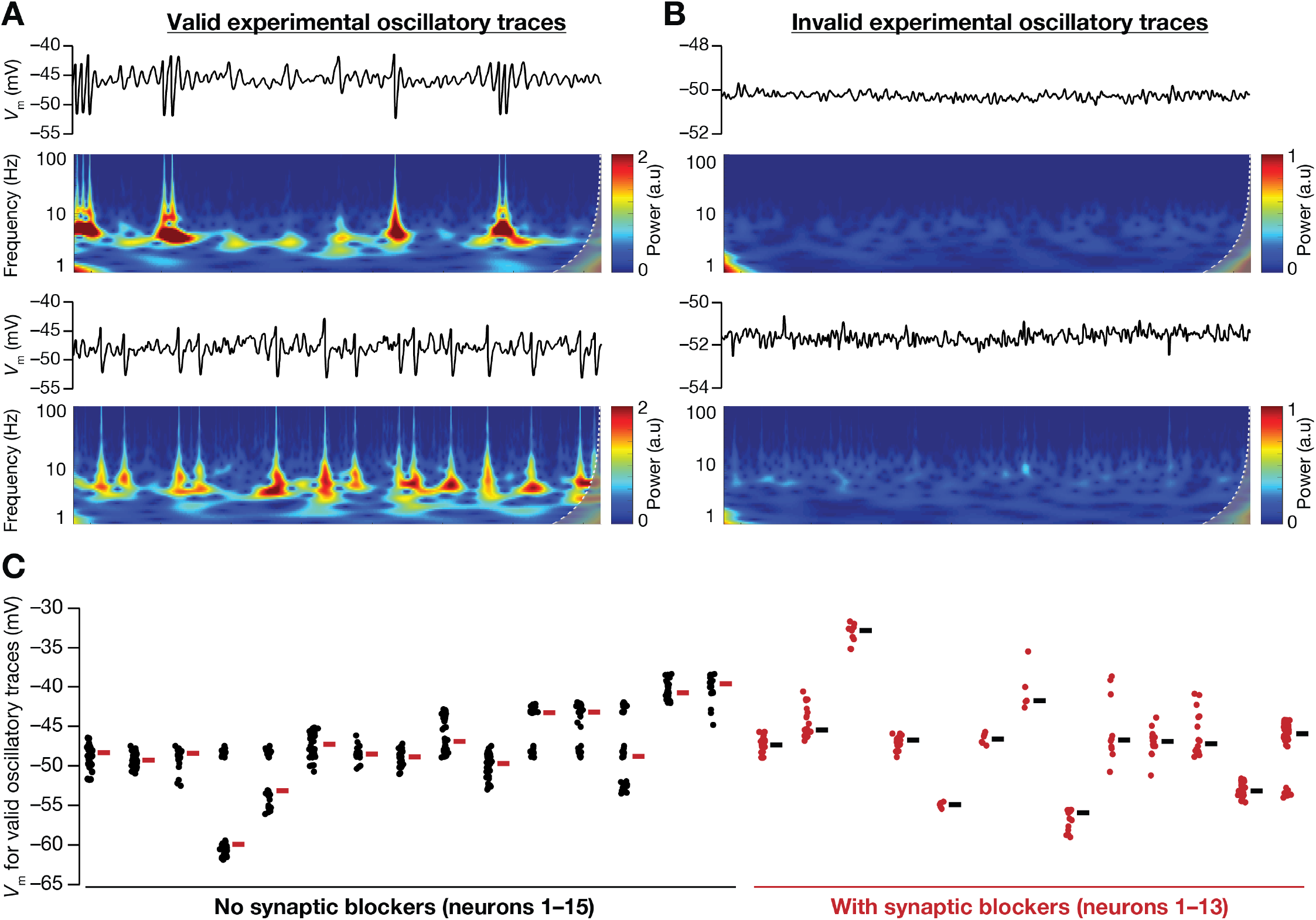
Electrophysiological recordings from rat LII MEC stellate cells showed valid intrinsic oscillatory traces at different peri-threshold membrane potential ranges. (A–B) Examples of valid (A) and invalid (B) oscillatory voltage traces (15 s long) and their respective wavelet transforms. Validation was performed using five measurements derived from the wavelet transform. (C) Mean membrane potentials where valid oscillatory traces were observed for each cell, recorded in the presence (red, *n* = 13) or absence (black, *n* =15) of synaptic receptor blockers.

**Figure 6:**
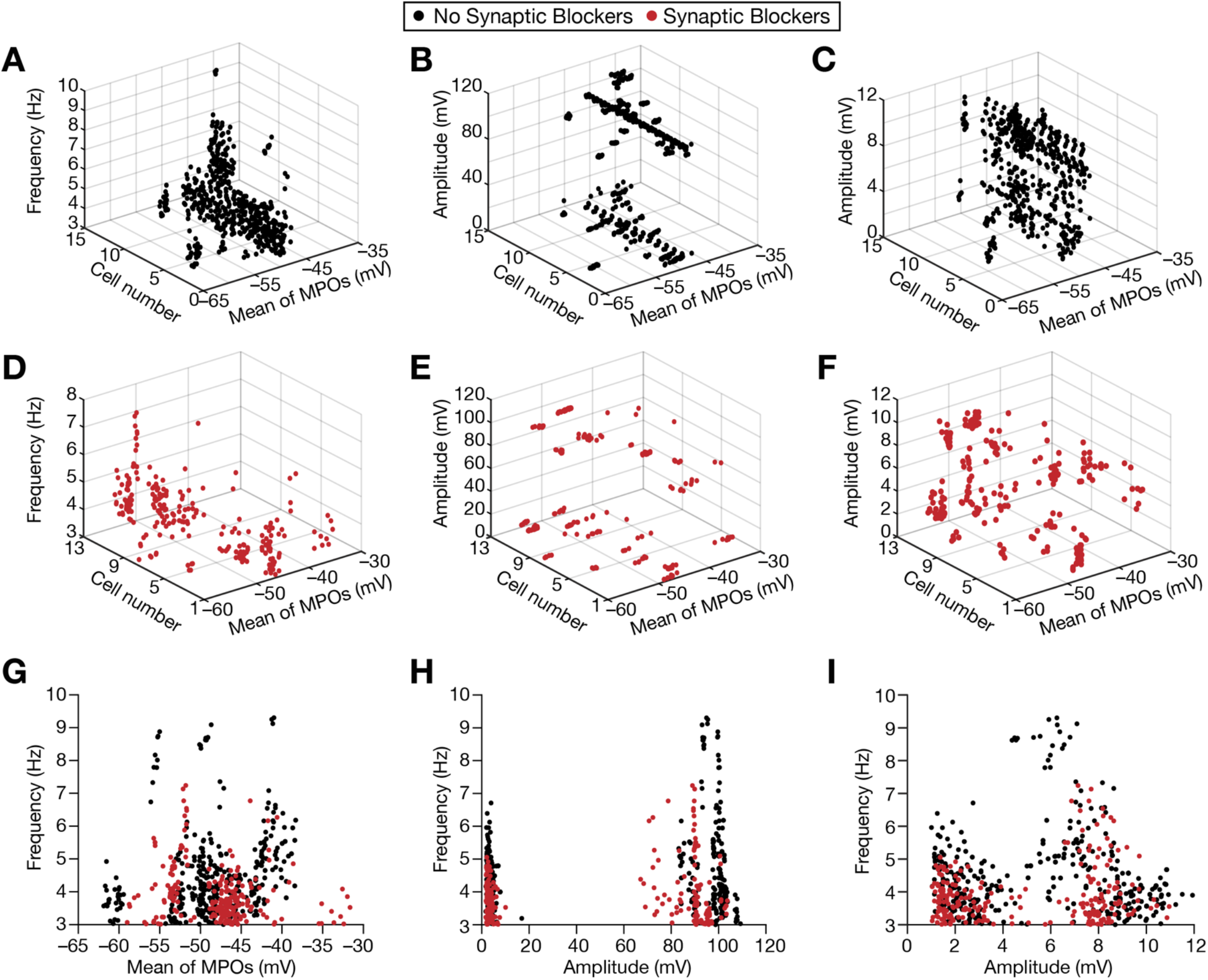
Heterogeneities in peri-threshold intrinsic oscillations observed in electrophysiological recordings from rat LII MEC stellate cells. (A–C) 3D-plots depicting conjunctive heterogeneities in frequency (A) or amplitude (B–C) and mean membrane potential where oscillatory traces were detected, across different cells. Amplitude plots are shown for unfiltered (B) traces to emphasize the manifestation of mixed-mode oscillations, or for median-filtered (C) traces to elucidate the amplitude and mean voltages of these oscillations when action potentials were removed. These traces were obtained in the absence of synaptic receptor blockers. (D–F) Same as panels (A–C), but for recordings obtained in the presence of synaptic receptor blockers. (G–I) Data from (A–F) depicted in 2D plots. Black and red dots represent measurements obtained without or with synaptic receptor blockers.

Together, analyses of peri-threshold activity patterns from our electrophysiological recordings confirmed that the five quantitative criteria that we set for validation of oscillatory traces (Fig. 4) were sufficient to capture the variable amplitude and irregular oscillatory patterns observed in MEC stellate cells (Fig. 1*I*, Figs. 5–6, Figs. S1–S2). These peri-threshold activity patterns were characteristically non-rhythmic, noisy, and of variable amplitude (Fig. 1*I*, Figs. 5–6, Figs. S1–S2), and were qualitatively similar to those observed with stochastic bifurcations in a simple nonlinear dynamical system (Fig. 2*C*, Fig. S4) and in stochastic models of MEC stellate cells (Fig. 3–4, Figs. S5–S6). Post-validation analyses of oscillatory traces unveiled pronounced cell-to-cell variability in the mean voltages where valid oscillations were detected, the amplitudes, and the frequencies of the valid oscillatory traces (Figs. 5–6).

### Manifestation of stochastic resonance in the detectability of valid oscillatory traces in the stochastic Hopf bifurcation system, but not with theta-filtered noise

We have developed quantitative metrics for validation of oscillatory traces (Fig. 4) and have confirmed that they can capture the variable amplitude and irregular oscillatory patterns observed in electrophysiological recordings from MEC stellate cells (Fig. 5–6). In addition, we had noted the qualitative similarities between valid subthreshold oscillatory traces from MEC stellate cells and those from the stochastic Hopf bifurcation. To further validate these metrics and to gain insights about the impact of noise on nonlinear dynamical systems showing bifurcations, we first applied these validation criteria for traces obtained from the stochastic Hopf bifurcation. As we were interested in the ability of stochasticity in stabilizing oscillatory traces, we assessed the dynamical system with the bifurcation parameter λ set at different values including those (λ ≤ 0) where they did not manifest stable limit cycles in the absence of noise. Specifically, when bifurcation parameter λ was greater than 0, the deterministic Hopf bifurcation manifested stable limit cycles in the temporal evolution (Fig. 7*A*; λ = 0.025; “No noise”), which would be detected as a valid oscillatory trace based on our criteria. However, with λ ≤ 0, the system manifested decaying spirals (Fig. 7*A*; λ =− 0.05, − 0.025, 0; “No noise”), which would not be identified as valid oscillatory traces.

**Figure 7:**
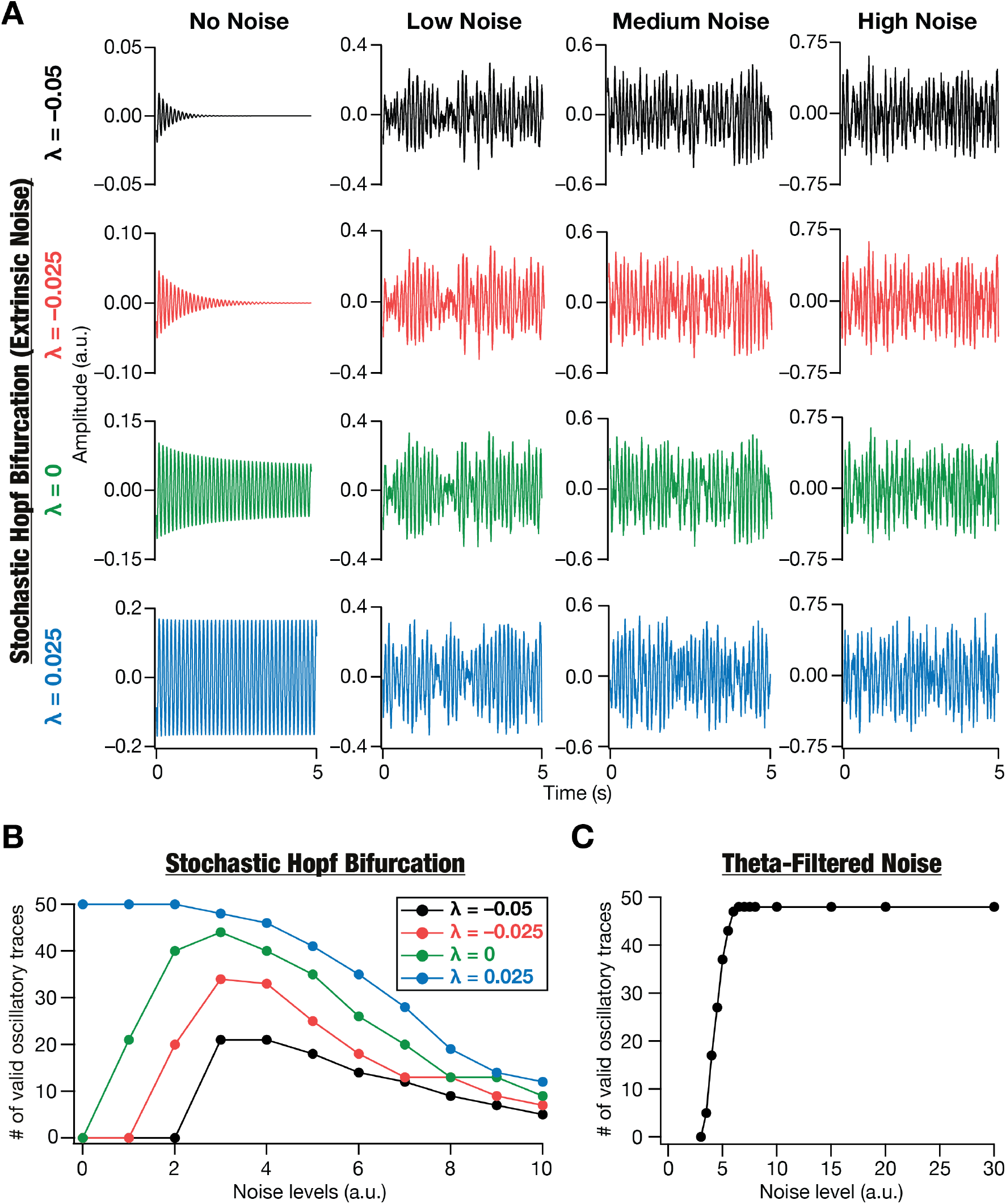
Stochastic resonance distinguishes filtered noise traces from noise-induced dynamics of a deterministic system manifesting a bifurcation yielding stable limit cycles. (A) Impact of different levels of Gaussian white noise (Low, Medium, High) on the dynamics of a deterministic Hopf bifurcation system, shown for different values of the bifurcation parameter λ. Note the emergence of stable oscillations in the deterministic system (“No Noise”) with λ > 0, and inward spirals with λ ≤ 0. (B) The number of valid oscillatory traces at different levels of noise for various values of the bifurcation parameter λ. Validation was performed on the outcomes of 50 trials for each value of λ at level of noise. Note the manifestation of stochastic resonance when λ ≤ 0: there is an optimal level of noise where the number of valid oscillatory traces is maximal, with the number falling on either side of this optimal level of noise. (C) The number of valid oscillatory traces at different levels of noise for various theta-filtered Gaussian white noise traces. Validation was performed on the outcomes of 50 trials for each level of noise. Notice a saturating monotonic increase in number of valid traces with increase in noise level, pointing to the absence of stochastic resonance with theta-filtered oscillatory traces.

**Figure 8:**
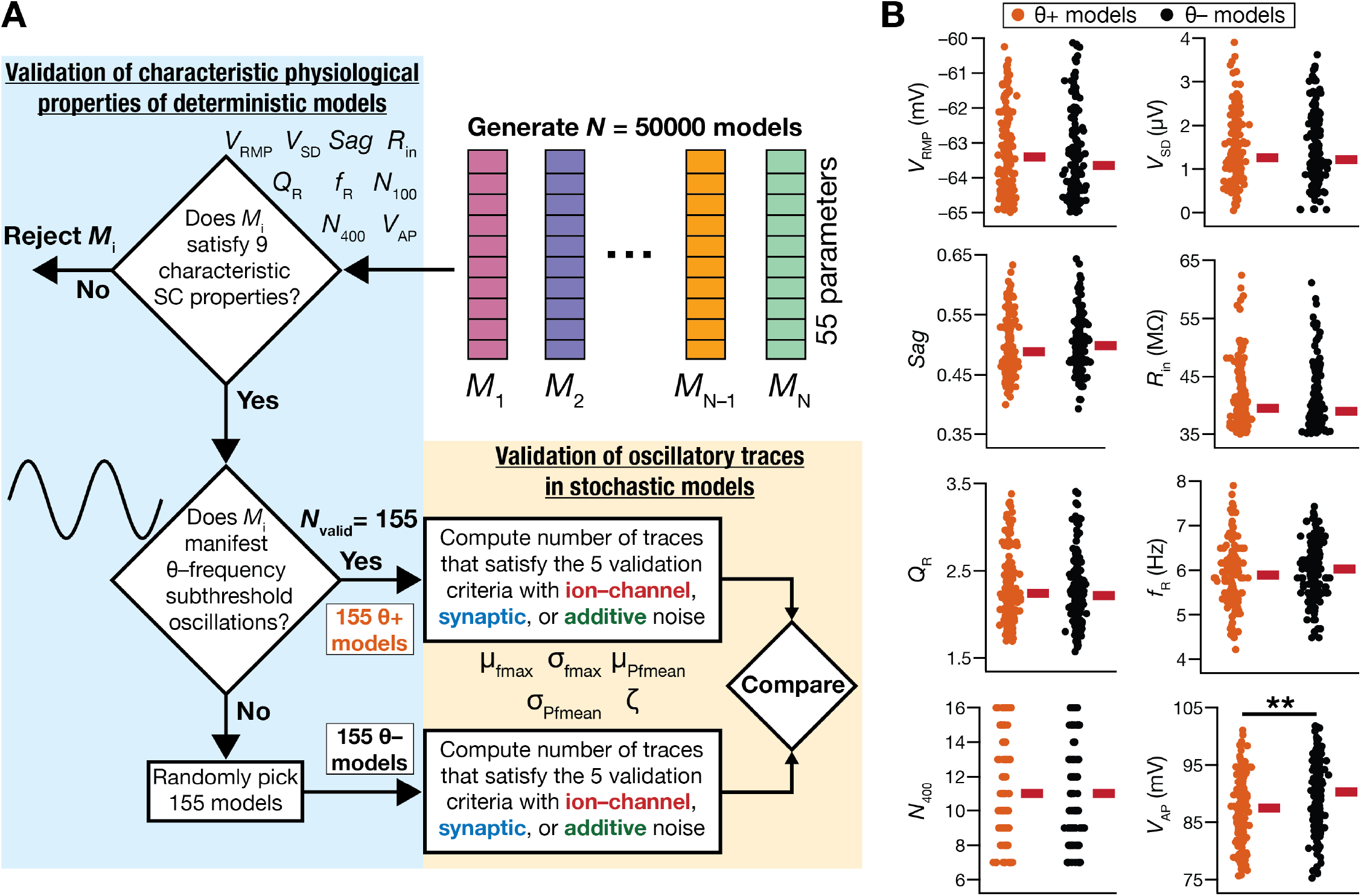
Definitions and measurement distributions of θ+ and θ– stellate cell models. (A) Flowchart representing the two-stage validation process, the first involving characteristic physiological properties of deterministic models (blue rectangle) and the second concerned with oscillatory traces in stochastic models (beige rectangle). Deterministic models that satisfied all ten characteristic physiological properties of entorhinal cortical stellate cells (Mittal and Narayanan, 2018) were defined as θ+ models. Models that satisfied 9 of the 10 characteristic physiological properties, but did not manifest deterministic, peri-threshold, theta-frequency oscillations were referred to as θ– models. (B) Distributions of the characteristic physiological properties of θ+ and θ– models. Shown are eight of the ten characteristic properties. As per the associated validation criterion, *N*_100_, the number of action potentials fired for a 100 pA current injection, was zero for all θ+ and θ– models (Mittal and Narayanan, 2018). The thick red line besides each measurement population represents the median for that population. *n* = 155 for both θ+ and θ– models. \*\**p*<0.005, Wilcoxon rank sum test. *p* value for *V*_AP_= 2.24 × 10^−4^. The *p* values for comparisons of all other measurements between θ+ and θ– models were >0.05.

We generated the temporal evolution traces from the Hopf bifurcation with different bifurcation parametric values and with different levels of stochasticity (Fig. 7*A*). We generated 50 traces for each level of noise at each value of λ, and subjected them to the validation criteria (Fig. 4) involving the five measurements derived from the spectrogram of these traces. As expected, validation was binary depending on the value of λ in the scenario with no noise, where there were no valid traces with λ ≤ 0 and all traces were valid when λ > 0. However, with increase in noise levels, we found that the decaying spirals in the deterministic system (when λ ≤ 0) switched to oscillatory patterns that manifested non-rhythmic and variable-amplitude oscillations in the stochastic system (Fig. 7*A*). As a consequence of these stochastic bifurcations introduced in the system with the introduction of noise, we found that the number of valid oscillatory traces increased with increased noise levels in scenarios where λ ≤ 0 (Fig. 7*B*). However, this increase in the detectability of valid oscillatory traces was not monotonic, and there was a fall in the detectability of oscillatory traces beyond a certain level of noise (Fig. 7*B*, λ ≤ 0). This phenomenon involving the manifestation of peak performance at an optimal level of noise, with performance falling on either side of this optimal level of noise, has been defined as stochastic resonance. In our case, the performance metric was the detectability of valid oscillatory traces or stabilization of unstable oscillations and there was indeed an optimal level of noise where the performance metric was at its peak (Fig. 7*B*, λ ≤ 0). With λ > 0, where stable oscillations were observed in the absence of noise, we observed a monotonic reduction in the number of valid oscillatory traces indicating the absence of stochastic resonance with λ > 0 (Fig. 7*B*).

As theta-filtered noise has also been postulated as a model for activity patterns in the MEC stellate cells, we assessed the impact of increasing noise intensity on theta filtered noise traces. Specifically, we subjected GWN to band-pass filtering in the theta frequency range and validated the resultant traces with the five metrics (Fig. 4) derived from their spectrograms. We repeated this for 50 trials with different seed values for generating the GWN and plotted the number of valid oscillatory traces as a function of GWN variance. Expectedly, and in striking contrast to stochastic bifurcations, filtered noise traces did not manifest stochastic resonance with increasing levels of noise, instead showing a saturating monotonic increase in the number of valid oscillatory traces with increasing noise levels (Fig. 7*C*). These analyses provide a clear quantitative demarcation between filtered noise and a system manifesting stochastic bifurcations in terms of the expression of stochastic resonance in response to increasing noise levels.

Together, these analyses further confirmed the reliability of the spectrogram-based metrics in capturing the stability of the oscillations in a stochastic nonlinear dynamical system. Importantly, these observations from a simple nonlinear dynamical system provided a key quantitative insight about stochastic bifurcations, unveiling the manifestation of stochastic resonance in these systems with specific reference to the role for noise in stabilizing oscillatory patterns. The expression of stochastic resonance indicates the presence of an optimal level of noise that facilitates stabilization of oscillatory patterns, with any other noise level (higher or lower) hampering optimal stabilization. In striking contrast, there was no stochastic resonance expressed with increasing noise levels in theta-filtered noise traces, and provides a quantitative handle to assess MEC stellate cells.

### The ability of stellate cell models to exhibit deterministic subthreshold oscillations translated to a greater number of valid oscillatory traces with the introduction of noise

Equipped with quantitative metrics for validation of oscillatory traces (Fig. 4), which we tested with electrophysiological recordings from rat MEC stellate cells (Figs. 5–6) and with oscillations in a nonlinear dynamical system (Fig. 7*A–B*), we returned to our original goal of quantitatively validating the 458,955 activity traces from the 155 stellate cell models. As mentioned earlier, visual inspection of traces provided qualitative indications for a stabilizing role of noise in the emergence of physiological activity patterns in MEC stellate cell models (Fig. 3). We computed the five quantitative metrics on all these traces that spanned three forms of noise (Fig. S7) to assess the ranges over which these five metrics were distributed. Of the 458,955 activity traces, we found a total of 129,309 oscillatory traces that satisfied the validation bounds on all five metrics. Although there were traces that failed each of the five criteria, the proportions of traces that failed the frequency stability and the minimum power criteria were the highest (Fig. S7). Importantly, we noted that with the introduction of noise, there were only a small proportion of traces (especially at low noise levels) that manifested inward or outward spirals, with most traces satisfying the spirality constraint (Fig. S7).

These distributions of the five metrics were derived from a population of 155 neurons that manifested deterministic subthreshold oscillations (referred to as θ+ models; Fig. 8*A*). Would these distributions be different if the deterministic models did not manifest subthreshold oscillations, but satisfied all the other characteristic physiological properties of MEC stellate cells (referred to as θ– models; Fig. 8*A*)? Would θ– models not manifest deterministic subthreshold oscillations switch to showing valid oscillatory traces with the introduction of these different forms of noise? To address these questions, we exploited the advantages of the multi-objective validation procedure in generating the deterministic MEC stellate cell models. Specifically, we randomly picked 155 deterministic models that satisfied 9 of the 10 characteristic physiological measurements that they were matched against, but did not manifest theta-frequency subthreshold oscillations (Fig. 8*A*). The number of these θ– models was set as 155 to match with the 155 θ+ models. We confirmed that the 9 other physiological measurements were comparable in θ+ and θ– models (Fig. 8*B*; the number of action potentials, *N*_100_, elicited by a 100-pA current injection was identically zero for all θ+ and θ– models). We then generated 10 independent trials of activity traces from each of these 155 θ– models at 21 different current levels, for distinct levels of the three different forms of noise. Together, this process yielded 458,955 activity traces from these θ– models, identical to the number of traces generated from θ+ models with the same process.

We computed the five quantitative metrics (for validation of oscillatory traces) on all traces from θ– models and compared their distributions with their counterparts from θ+ models (Fig. 8*A*). Overall, there was a three-fold lesser number of valid oscillatory traces generated from θ– models (42,477 out of 458,955) compared to their θ+ counterparts (129,309 out of 458,955). In addition, we compared the distributions of each of the five quantitative metrics across θ+ and θ– models, for each of the different forms of noise (Fig. S8). Although the distributions for μ_fmax_ and ζ were comparable, especially given the validation bounds on these metrics, there were considerable differences in the other three metrics derived from θ+ and θ– models (Fig. S8). Specifically, there was a larger fraction of θ– model traces (compared to θ+ model traces) that failed the frequency stability criterion because σ_fmax_ was greater than 1 Hz for a larger fraction of traces (Fig. S8). Although the power stability criterion on σ_Pfmean_ was satisfied by a larger proportion of θ– model traces (compared to θ+ model traces), this was simply because the overall power of oscillations was lower in traces from θ– models (Fig. S8). As a consequence of the dominantly low-power distribution of θ– model traces, there were more validation failures for θ– model traces compared to their θ+ counterparts owing to the validation requirement that μ_Pfmean_ > 0.5.

Together, these analyses demonstrated that the existence of a bifurcation state that manifested subthreshold oscillations in the deterministic model translated to enhanced propensity of the model to express peri-threshold oscillatory activity in the presence of different forms of noise. The absence of a bifurcation state that manifested subthreshold oscillations in the deterministic model (θ– models) resulted in peri-threshold activity patterns that were of lower power with unstable peak frequencies across time.

### Stochastic resonance in the emergence of peri-threshold oscillatory activity in MEC stellate cell models

Our analyses of peri-threshold activity traces, at the population level involving all noise levels at each form of noise, showed the presence of valid oscillatory patterns in both θ+ and θ– models. How does the level of noise influence the number of valid oscillatory traces obtained from the population of θ+ and θ– models? We had earlier demonstrated the expression of stochastic resonance with increased levels of noise in a system manifesting stochastic bifurcation, but not with filtered noise (Fig. 7*B–C*). The expression of stochastic resonance in the detectability of valid oscillatory traces from MEC stellate cell models would strengthen our central hypothesis that peri-threshold activity patterns in MEC stellate cells are consistent with those elicited by stochastic bifurcations in a heterogeneous neuronal population. To assess this, we plotted the number of valid oscillatory traces observed with each level of noise across the three forms of noise applied to the heterogeneous population of θ+ and θ– models (Fig. 9). Strikingly, we found the manifestation of stochastic resonance in the detectability of valid oscillatory traces obtained from both θ+ and θ– model populations for forms of noises, spanning different values of injected current (Fig. 9*A–C*). Specifically, there was an optimal level of noise, for each of ion-channel, synaptic, and additive noise, where the models manifested the highest number of valid oscillatory traces; there were lesser number of valid traces at noise levels on either side of this optimal noise level (Fig. 9*A–C*).

**Figure 9:**
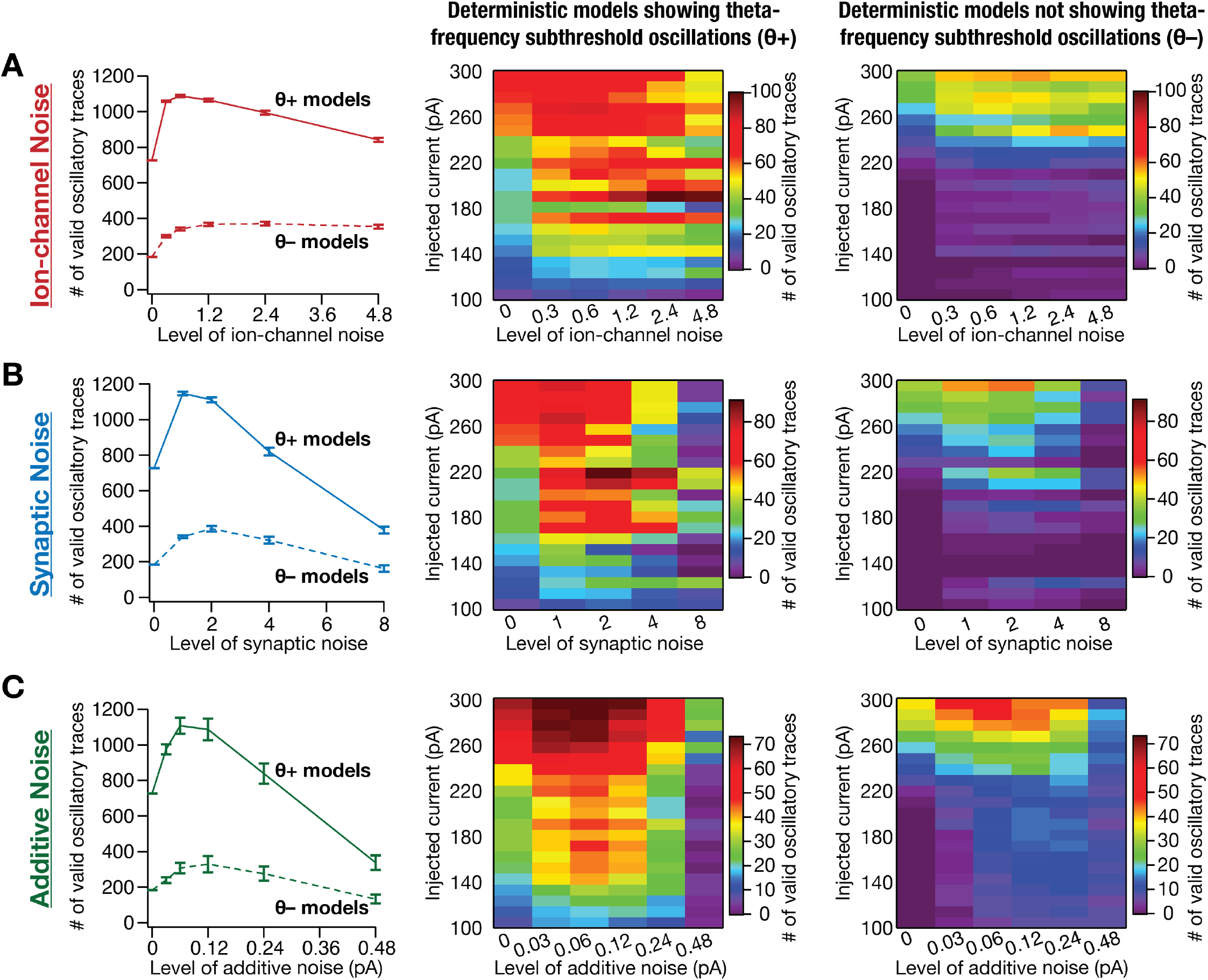
Stochastic resonance in the emergence of peri-threshold oscillations in a heterogeneous population of MEC stellate cell models. (A–C) *Left,* Mean and SEM of number of valid oscillatory traces from all θ+ (*n*_θ+_ = 155) and θ– (*n*_θ–_ = 155) model neurons. *Middle,* Number of valid oscillatory traces spanning all 21 current injection (*I*_inj_) values across different levels of noise across all θ+ model neurons. *Right,* Same as the middle panel, but for θ– model neurons. The number of valid oscillatory traces is plotted as mean and SEM spanning 10 independent trials for each level of the three forms of noise: (A) ion-channel noise, (B) synaptic noise, or (C) additive noise, for all θ+ and θ– models. Total number of valid traces: ion-channel noise, θ+ (50,509/162,750) and θ– (17,320/162,750); synaptic noise, θ+ (34,588/130,200) and θ– (12,112/130,200); additive noise, θ+ (43,485/162,750) and θ– (12,861/162,750); deterministic models (0 noise), θ+ (727/3,255) and θ– (184/3,255).

Although traces from θ– models manifested stochastic resonance, the total number of valid traces at each noise level was considerably lower compared to those from θ+ models, across all noise levels and all forms of noise. Importantly, in the tested range of noise levels with synaptic and additive noise, as noise crossed a certain threshold, the detectability of valid oscillatory traces was lower compared to the deterministic scenario where there was no noise (Fig. 9*B–C*). The number of valid oscillatory traces for deterministic (no noise) θ– models was non-zero because θ– models were picked based on the absence of subthreshold theta-frequency oscillations, implying a requirement that there are no spikes. However, the five criteria employed for validating oscillatory traces were designed to validate mixed-mode oscillations, that manifest spikes, as well. The presence of mixed-mode oscillations in these traces is also evident from the observation that valid oscillatory traces in θ– models were largely confined to higher current injection values between 220–300 pA (Fig. 9*A–C*, right panels).

### Stochastic resonance in the emergence of peri-threshold oscillatory activity in individual MEC stellate cell models

Our analyses of the manifestation of stochastic resonance in observing valid peri-threshold oscillations in MEC stellate cell models were at the population level, involving traces from all models (Fig. 9). Thus, these analyses did not address the question of whether stochastic resonance in observing valid peri-threshold oscillations manifested at the single-neuron level. Specifically, these analyses did not provide evidence for the existence of an optimal noise level of noise that increases the probability of observing peri-threshold oscillations in individual neurons. To address this, we performed two sets of analyses assessing the number of valid traces in individual models for different forms and levels of noise (Fig. 10*A–B*).

**Figure 10:**
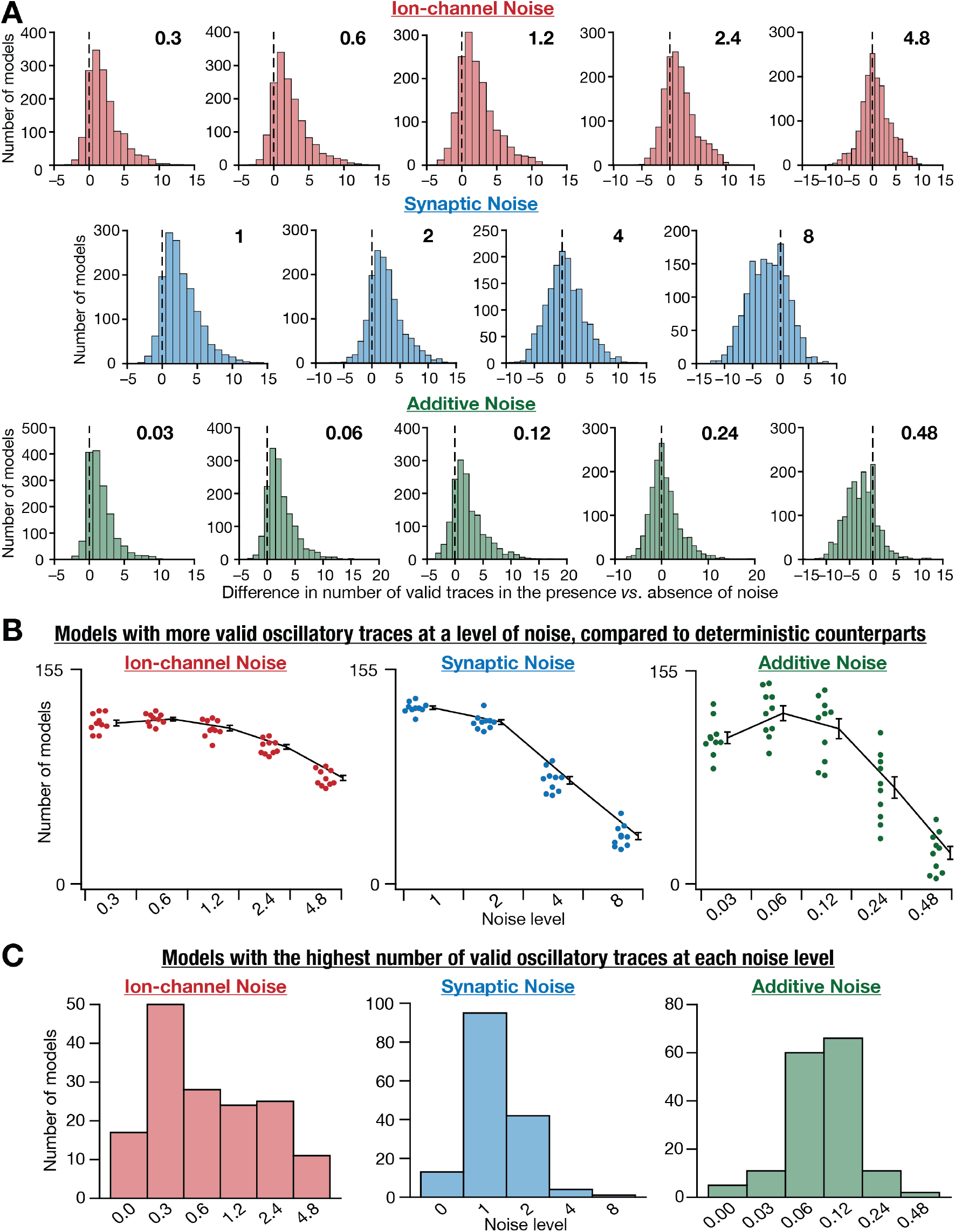
Stochastic resonance in the emergence of peri-threshold oscillations in individual MEC stellate cell models. (A) Histograms of the differences in the number of valid oscillatory traces in the presence *vs*. the absence of noise. Positive differences indicate a beneficial impact of noise on the number of valid traces. Histograms are shown for each level of noise (different columns) for all three forms of noise (different rows). These histograms indicated pooled data from 10 different trials for each level of all forms of noise, across all 155 θ+ models. (B) Summary data from derived from panel A, showing the number of models that yield more valid oscillatory traces in the presence of noise (compared to deterministic models with no noise), at each of the different levels of all forms of noise. The individual data points represent each of the 10 trials for a given level of noise, and the summary statistics are represented as mean and SEM across trials. (C) The number of neuronal models (out of the maximum possible 155 θ+ models) exhibiting the highest number of valid oscillatory traces at each level of the three forms of noise. Noise level 0 indicates deterministic models. The maximum for each model was computed by considering the number of valid oscillatory traces in the model for all levels of a specific form of noise.

As each model was assessed at 21 different current values, 21 is the maximum number of valid oscillatory traces from an individual model for a given noise level of a specific form of noise, a bound that holds for the zero-noise deterministic scenario as well. Let 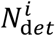 and 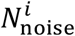 be the number of valid oscillatory traces in model *i* in the absence (deterministic) or presence of noise, respectively. We computed the difference 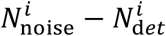 for all θ+ models (1 ≤ *i* ≤ 155), for 10 trials at each level of the three forms of noise and plotted the distributions of these differences spanning all models and trials (Fig. 10*A*). These distributions could theoretically span the range from –21 to +21, with positive values indicating that noise enhanced the number of valid oscillatory traces and negative values implying a deleterious impact of noise on this number. We then counted the number of model neurons that showed a greater number of valid oscillatory traces upon introduction of noise (Fig. 10*B*), in comparison to their deterministic counterpart (*i.e.*, a positive value of 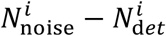). We found that there were significant proportion of models manifested more oscillatory traces in the presence of noise, across all 10 trials, indicating a role for noise in stabilizing oscillation (Fig. 10*B*). In addition, the number of neurons with positive values for 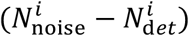 was the highest at optimal noise levels, with the number falling on either side, together providing evidence for the expression of stochastic resonance in the expression of peri-threshold oscillations at the single-neuron level (Fig. 10*A–B*).

Second, we noted down the noise level at which each individual model showed the highest number of valid oscillatory traces, with the maximum computed across all levels of a given form of noise for that specific model. We noted that different models showed the maximal number of valid oscillatory traces for different levels of noise, thus demonstrating heterogeneities in the specific value of the optimal level of noise across models (Fig. 10*C*). We binned all 155 θ+ models based on the specific noise level that they showed the maximum number of valid oscillatory traces (Fig. 10*C*). We observed that very few models showed maximal number of valid oscillatory traces in the absence of noise, thus emphasizing a positive role for noise in the manifestation of peri-threshold oscillations (Fig. 10*C*). Thus, these analyses unveiled neuron-to-neuron heterogeneities in the optimal level of noise required for the optimal manifestation of peri-threshold oscillations in individual neurons, providing further lines of quantitative evidence for the expression of stochastic resonance in the manifestation of peri-threshold oscillations.

As a final line of evidence for the manifestation of stochastic resonance in the expression of peri-threshold oscillations at the level of individual neurons, we normalized the number of valid traces in each model with reference to the maximum number of valid traces spanning all noise levels of a specific form of noise. This allowed us to account for the heterogeneities across different models, whereby each model manifested different numbers of maximal valid oscillatory traces at disparate levels of noise (Fig. 10*C*). By normalizing the number of oscillatory traces across all noise levels for individual models, we have a plot for each model which is 1 at the specific noise level where it attained its maximal value and would be ≤ 1 at other noise levels. This normalization ensured that there was no domination by models with a greater number of valid traces. We now took these plots for each of the 155 θ+ models and plotted their mean as a function of noise level for each form of noise (Fig. S9*A*). Consistent with our earlier conclusions (Fig. 10), we found that the average fraction of valid traces in individual models was highest at a specific optimal level of noise, for each of the three forms of noise (Fig. S9*A*). Considering the heterogeneities across models in terms of the maximal number of valid oscillatory traces and the optimal noise level at which they manifested this maximal number, we asked if the frequency of these oscillations were different across different noise levels across the three different forms of noise (Fig. S9*B*). Although there were heterogeneities in the frequency of oscillations across traces, we found the range of oscillations and summary statistics to be comparable across noise levels for all forms of noise (Fig. S9*B*).

We had set forth with the assessment of stochastic resonance in this heterogeneous model population guided by the manifestation of stochastic resonance in the simple nonlinear dynamical system endowed with stochastic bifurcations (Fig. 7*A–B*), but not with filtered noise (Fig. 7*C*). We reasoned that the expression of stochastic resonance in MEC stellate cells models would constitute a line of evidence for stochastic bifurcations as an underlying mechanism for activity patterns in these models. Specifically, if the activity patterns were merely filtered noise, increase in noise levels should have monotonically increased the power in those frequencies, yielding more stable oscillations in the filtered frequency range (Fig. 7*C*). On the other hand, if the LII MEC models manifested stochastic resonance, we reasoned that such a scenario would be consistent with the expression of stochastic bifurcations in a nonlinear dynamical system (Fig. 7*A–B*). Our analyses (Figs. 9–10) demonstrated that the introduction of noise in a heterogeneous population of deterministic stellate cells manifesting bifurcations allowed for the stabilization of peri-threshold oscillations with an optimal level of noise. These observations provided clear lines of evidence for the expression of stochastic resonance in the detectability of oscillations with increasing noise power, providing evidence against the possibility that activity patterns in MEC stellate cells were merely filtered noise patterns.

## DISCUSSION

The principal goal of this study was to identify a theoretical framework that best describes the peri-threshold activity patterns observed in MEC stellate cells (Alonso and Llinas, 1989; Alonso and Klink, 1993), towards resolving the disparate characterizations of these activity patterns. Prior studies have proposed modeling these activity patterns to be periodic oscillations, emergent through interactions between specific ion channels (Klink and Alonso, 1993; White et al., 1993; Dickson et al., 2000b; Fransen et al., 2004) or as noise filtered in the theta frequency range (Dodson et al., 2011). It is evident from recordings from MEC stellate cells that the simple periodic oscillator framework fails because activity patterns from these neurons manifest noisy oscillations with variable frequency, phase, and amplitude. In addition, there are lines of evidence for the absence of such oscillatory patterns in intracellular whole cell *in vivo* recordings from these stellate cells, conditions where they are faced with high intrinsic and synaptic noise levels (Schmidt-Hieber and Hausser, 2013). Under the filtered noise framework, these conditions should instead have yielded stable oscillations (Fig. 7*C*), thus providing evidence against the filtered noise for explaining these activity patterns. In this study, using a combination of theoretical, computational, and electrophysiological methods coupled with rigorous quantitative analyses, we argue for heterogeneous stochastic bifurcations as a unifying framework that explains all aspects of these peri-threshold activity patterns.

### Stochastic bifurcations and biological systems

There are several examples within the dynamical systems literature showing a considerable impact of stochasticity on the bifurcations, including shift in the bifurcation points, introduction of secondary bifurcations, and manifestation of bistability (Sastry and Hijab, 1981; Namachchivaya and Ariaratnam, 1987b, a; Sri Namachchivaya, 1988, 1990; Colonius and Kliemann, 1994; Juel et al., 1997; Aumaître et al., 2007). The stochastic fluctuations in microscopic ion-channel dynamics result in considerable differences in macroscopic properties, and need to be specifically accounted for if they were to be matched with biological neuronal properties (White et al., 1998; Dorval and White, 2005; Dudman and Nolan, 2009; Cannon et al., 2010; Dodson et al., 2011). The presence of stochastic bifurcations and stochastic resonance in biological systems has been studied not just at the cellular scale, but across all scales of biological systems from the perspective of bistability as well as in terms of stabilizing oscillatory properties (Bulsara et al., 1991; Longtin et al., 1991; Chialvo and Apkarian, 1993; Douglass et al., 1993; Longtin, 1993; Braun et al., 1994; Collins et al., 1995; Cordo et al., 1996; Gluckman et al., 1996; Levin and Miller, 1996; Simonotto et al., 1997; Russell et al., 1999; Mori and Kai, 2002; Priplata et al., 2003; Losick and Desplan, 2008; Raj and van Oudenaarden, 2008; Song et al., 2010; Tyson and Novak, 2010; Ishikawa, 2015; Bashkirtseva et al., 2016; Fu et al., 2018; Alon, 2019). Thus, in assessing bifurcations in neural activity patterns, it is important that they are not mapped onto deterministic bifurcations emergent from macroscopic models of ion channel function, but as stochastic bifurcations that account for fluctuations in microscopic components. Deterministic models that deal with macroscopic dynamics in a deterministic fashion run the risk of not matching biological properties under different contexts with distinct forms of extrinsic and intrinsic perturbations.

### Peri-threshold activity patterns in MEC stellate cells as emergent dynamics in a heterogeneous population of neurons manifesting stochastic bifurcations

We demonstrate that activity patterns in MEC stellate cells can be explained by a theoretical framework that considers them as emergent dynamics of activity in a heterogeneous population of neurons manifesting stochastic bifurcations. First, a simple nonlinear dynamical system showing a deterministic bifurcation that results in stable limit cycles switched to showing variable amplitude, arhythmic oscillatory patterns upon the introduction of noise (Fig. 2*C*). The nature of oscillations in this simple system manifesting stochastic bifurcations motivated us to introduce noise into deterministic models of MEC stellate cells showing bifurcations (from rest to subthreshold oscillations to action potential firing). We used a heterogeneous population of conductance-based LII stellate cell models and demonstrated that the introduction of three different forms of noise stabilized oscillatory patterns in these model neurons (Fig. 3, Figs. S5– S6). Specifically, we show that the activity patterns elicited by these stochastic model neurons were qualitatively similar to LII stellate cells (irregular, non-rhythmic, variable amplitude, and noisy), although their deterministic counterparts manifested unphysiological patterns such as decaying or expanding oscillations (Fig. 3, Figs. S5–S6). Importantly, deterministic models manifesting regular subthreshold oscillations switched to showing non-rhythmic, variable amplitude oscillations, and models manifesting regular spiking switched to mixed-mode oscillations (spikes riding on top of subthreshold oscillations, with certain cycles skipped) with the introduction of stochasticity. Thus, model MEC stellate cells manifested stochastic bifurcations in the presence of noise, switching from rest to subthreshold oscillations to mixed-mode oscillations to regular action potential firing, with injected current acting as the bifurcation parameter. These stochastic bifurcations were heterogeneous and manifested considerable variability in a manner that was dependent on the specific model and on the level and form of noise.

To quantitatively assess stabilization of these oscillatory patterns in the presence of noise, we defined five different metrics from the spectrogram of unfiltered voltage traces for detecting stable theta-frequency oscillations (Fig. 4). We confirmed with peri-threshold oscillations in electrophysiological recordings that these metrics were sufficient to capture the variable amplitude and irregular oscillatory patterns observed in MEC stellate cells (Fig. 5–6). We also found these metrics to reliably capture stability of oscillations in a stochastic Hopf bifurcation system (Fig. 7*A*). Validation of traces from the stochastic Hopf bifurcation system provided a key insight about the emergence of stochastic resonance with increasing levels of noise (Fig. 7*B*). The expression of stochastic resonance indicates the presence of an optimal level of noise that facilitates stabilization of oscillatory patterns, with any other noise level (higher or lower) hampering optimal stabilization. In striking contrast, there was no stochastic resonance expressed with increasing noise levels in theta-filtered noise traces, thus providing us a quantitative handle to ask if activity patterns in MEC stellate cells are best characterized by stochastic bifurcations, and not by theta-filtered noise (Fig. 7*B–C*).

We assessed two populations of heterogeneous deterministic MEC stellate cell models based on their ability to manifest a bifurcation state involving subthreshold theta-frequency oscillations (Fig. 8). The presence of a bifurcation state that manifested subthreshold oscillations in the deterministic model translated to enhanced propensity of the model to express peri-threshold oscillatory activity in the presence of noise (Fig. 9). We validated traces with different forms of noise and unveiled the expression of stochastic resonance in the emergence of peri-threshold oscillations in MEC stellate cell models (Figs. 9–10). Importantly, there are lines of evidence for the expression of stochastic resonance in MEC stellate cells from *in vitro* experiments. Specifically, it has been shown that additional ion-channel or synaptic noise, introduced under *in vitro* conditions using a dynamic clamp setup, play a critical role in the emergence of peri-threshold oscillations (White et al., 1998; Dorval and White, 2005; Fernandez and White, 2008; Fernandez et al., 2015). Our analyses provided quantitative evidence for the manifestation of heterogeneous stochastic bifurcations in MEC stellate cell population, and against the possibility that peri-threshold activity patterns were merely filtered noise.

Heterogeneous stochastic bifurcations as the theoretical framework for explaining intrinsic patterns in MEC stellate cells implies state-dependence of both synaptic integration and the specific types of patterns emerging from these neurons. For instance, the expression of stochastic resonance provides a quantitative explanation for the absence of peri-threshold oscillations in intracellular *in vivo* recordings from stellate cells (Schmidt-Hieber and Hausser, 2013), where the noise levels are high beyond the optimal level of noise (Figs. 9–10) required for their emergence (Fernandez and White, 2008). Within this framework, under *in vitro* conditions, even with the blockade of synaptic receptors and the absence of additional external noise (Fig. 1, Figs. 5–6, Figs. S1–S2), the intrinsic ion-channel noise is sufficient to elicit peri-threshold oscillations with variable amplitude and frequency. The heterogeneous stochastic bifurcations framework also implies that synaptic integration in MEC stellate cells depends on the composition and properties of ion channels in each cell, the form and level of noise encountered by individual compartments, and the specific timings and location of synaptic inputs (White et al., 1998; Dorval and White, 2005; Fernandez and White, 2008; Fernandez et al., 2015).

As stochastic bifurcations that are central to this framework are mediated by stochastic ion channels, differences in these oscillatory patterns with the blockade of different ion channels (including HCN, persistent sodium, *M*-type K^+^) are also readily explained within this framework (Mittal and Narayanan, 2018). Therefore, activity-dependent plasticity, neuromodulation, or channelopathies that affect ion channels that are active in the peri-threshold range could alter these activity patterns. Further, the neuron-to-neuron variability manifested by peri-threshold oscillatory patterns are driven by the stochasticity and the underlying heterogeneities in the deterministic neuronal population. Specifically, the constellation of characteristic physiological properties of MEC stellate cells could be achieved through disparate combinations of ion channels and other parameters, implying the manifestation of degeneracy (Mittal and Narayanan, 2018). This implied heterogeneities in the expression profile of individual ion channels in each model, resulting in different kinds of activity patterns (regular subthreshold oscillations, regular spiking, no oscillations, decaying oscillations, expanding oscillations), and differential interactions with noise. The strength and type of noise added an additional layer of variability in how these oscillatory patterns manifested, together introducing pronounced neuron-to-neuron variability observed in electrophysiological recordings. These variabilities also emphasize the need to use a heterogeneous population of neuronal models with different levels and forms of noise in assessing these activity patterns, as there would be considerable biases and misinterpretations with the use of a single deterministic hand-tuned model.

Together, these analyses provide clear lines of evidence for peri-threshold activity patterns in MEC stellate cells to be consistent with the manifestation of heterogenous stochastic bifurcations in these cells, explaining all signature properties of these activity patterns spanning different experimental conditions.

### Future directions

The computational complexity of assessing heterogeneous stochastic bifurcations in our neuronal model population was enormous, requiring each of the multiple models to be assessed at different current injections with different levels and forms of noise. Therefore, we had resorted to the use of single compartmental conductance-based models, accounting for all the channel kinetics and intrinsic properties of the stellate cells. However, to assess the impact of stochasticity and heterogeneity on neuronal activity patterns, it is essential that electrophysiological studies characterize dendritic ion-channel and intrinsic properties across the arbor of MEC stellate cells. A heterogeneous population of morphologically realistic models could then be built to assess the impact of stochasticity (ion-channel and synaptic noise) and heterogeneities (in biophysical, synaptic, and morphological properties) on location-dependent peri-threshold oscillatory patterns and synaptic integration. Such models would also enable assessment of context-dependence of stellate cell physiology, with levels and forms of noise, activity-dependent plasticity of channels and receptors, neuromodulation, and pathological channelopathies driving the context for physiological changes. The morphologically realistic models also would enable the introduction of balanced synaptic noise in a location-dependent manner, thus providing a more detailed assessment of the impact of synaptic noise and balance therein on intrinsic activity patterns.

In introducing ion-channel noise, we introduced stochasticity solely in the persistent Na^+^ channel. While it is possible to introduce stochasticity in all the channels, but we have focused on persistent Na^+^ channel for three reasons. First, in studies involving electrophysiological recordings and computational modeling, it has been shown that blockade of persistent Na^+^ channels result in complete loss of peri-threshold oscillations (Alonso and Llinas, 1989; Klink and Alonso, 1993; Boehlen et al., 2013; Mittal and Narayanan, 2018). Second, previous studies have reported the role of stochastic persistent Na^+^ channels in emergence of canard structure which is important for mixed-mode oscillations, also pointing toward role of stochasticity in stabilization of oscillations (Dorval and White, 2005). Lastly, we have 9 active and leak conductances in the model neurons with each model constructed with a unique set of parameters. The computational complexity in accounting for stochasticity in each channel was enormous, especially given the number of models, trials, and noise configurations required for unbiased evaluation. Future studies could employ detailed stochastic kinetic models for each ion channel to assess the impact of stochasticity and heterogeneities on peri-threshold activity patterns.

One of the important electrophysiologically testable predictions from our study is the expression of stochastic resonance in the emergence of peri-threshold oscillations in MEC stellate cells. While it is impossible to achieve zero-noise conditions in biological neurons endowed with intrinsically stochastic ion channels, interventional experiments involving introduction of additional noise (White et al., 1998; Dorval and White, 2005; Fernandez and White, 2008; Fernandez et al., 2015) or suppressing high synaptic noise conditions could be performed. Specifically, peri-threshold activity patterns could be recorded from MEC stellate cells across the dorso-ventral axis under *in vivo* conditions with different amplitudes of current injection, in the presence and absence of synaptic receptor blockers. These activity patterns could then be subjected to our validation metrics to address questions of whether oscillations emerge when the high synaptic noise is suppressed, and if there are differences in dorsal *vs*. ventral oscillatory patterns *in vivo*. If there are more valid oscillatory traces in the presence of synaptic blockers, that would provide direct electrophysiological evidence for the expression of stochastic resonance. Similar *in vivo* electrophysiological experiments could be performed with blockers for HCN, persistent sodium, or *M*-type K^+^ channels to assess the number of valid oscillatory traces in their presence *vs.* absence. These experiments would provide evidence for the role of these ion channels in mediating the stochastic bifurcations that are postulated here to mediate peri-threshold oscillations.

## ACKNOWLEDGMENTS

The authors thank members of the cellular neurophysiology laboratory for helpful discussions and for comments on a draft of this manuscript. The authors thank Dr. Arvind Kumar and Dr. Matthew Nolan for helpful discussions. This work was supported by the Wellcome Trust-DBT India Alliance (Senior fellowship to R. N.; IA/S/16/2/502727), the Department of Biotechnology through the DBT-IISc partnership program (R. N.), the Revati and Satya Nadham Atluri Chair Professorship (R. N.), and the Ministry of Human Resource Development (R. N. & D. M.).

## AUTHOR CONTRIBUTIONS

D. M. and R. N. designed experiments; D. M. performed experiments and carried out data analysis; D. M. and R. N. co-wrote the paper.

## DATA AVAILABILITY

All data needed to evaluate the conclusions in the paper are present in the paper and/or the Supplementary Materials.

