## Supplementary Figures S1-S9; Supplementary Table S1-S2 for "Stochastic resonance and bifurcations in a heterogeneous neuronal population explain intrinsic oscillatory patterns in entorhinal cortical stellate cells"

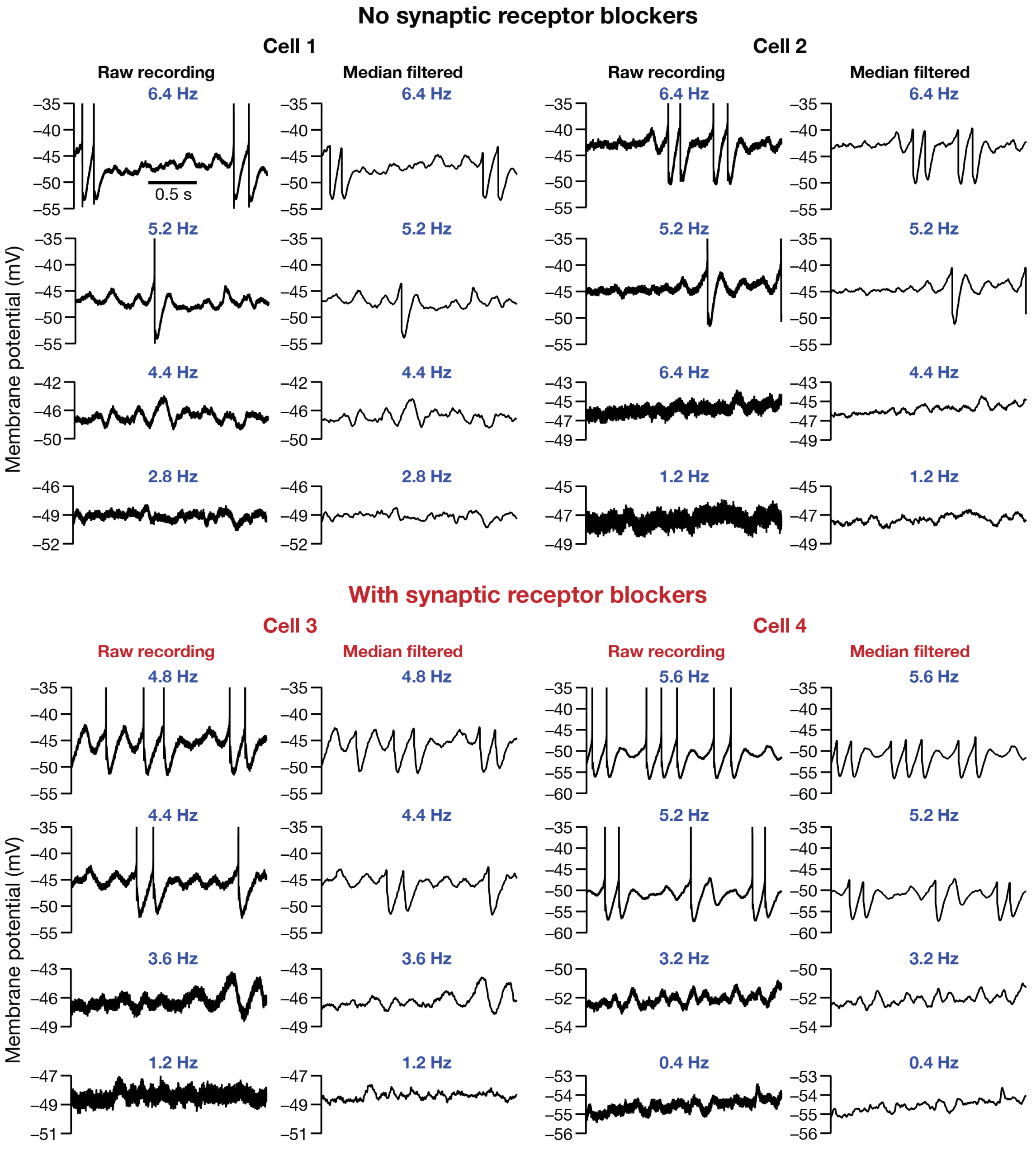

**Supplementary Figure S1: Example voltage traces from electrophysiologically recorded LII MEC stellate cells using the short oscillation protocol.** Each column depicts four sets of traces recorded at different voltage levels from a single neuron. For each cell, the raw recordings (*Left*) along with the median filtered version of same trace (*Right*) are shown. Cells 1–2 were recorded in the absence of synaptic blockers (black), whereas Cells 3–4 were recorded in the presence of synaptic blockers (red). Note that when spikes occurred, they were truncated to –35 mV for emphasizing the subthreshold dynamics.

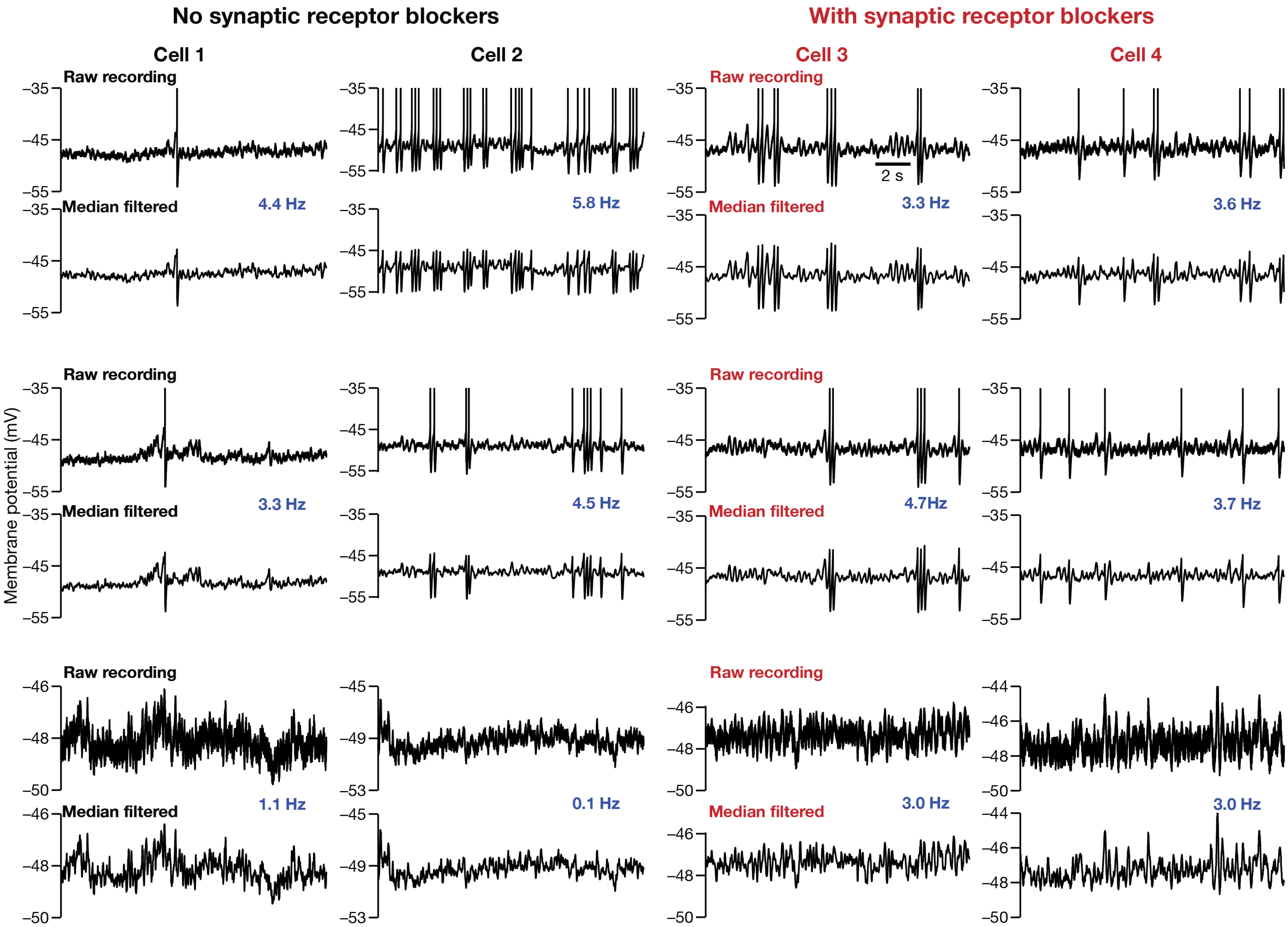

**Supplementary Figure S2: Example voltage traces from electrophysiologically recorded LII MEC stellate cells using the short oscillation protocol.** Each column depicts three sets of traces recorded at different voltage levels from a single neuron. For each cell, the raw recordings (*Top*) along with the median filtered version of same trace (*Bottom*) are shown. Cells 1–2 were recorded in the absence of synaptic blockers (black), whereas Cells 3–4 were recorded in the presence of synaptic blockers (red). Note that when spikes occurred, they were truncated to –35 mV to emphasize subthreshold dynamics.

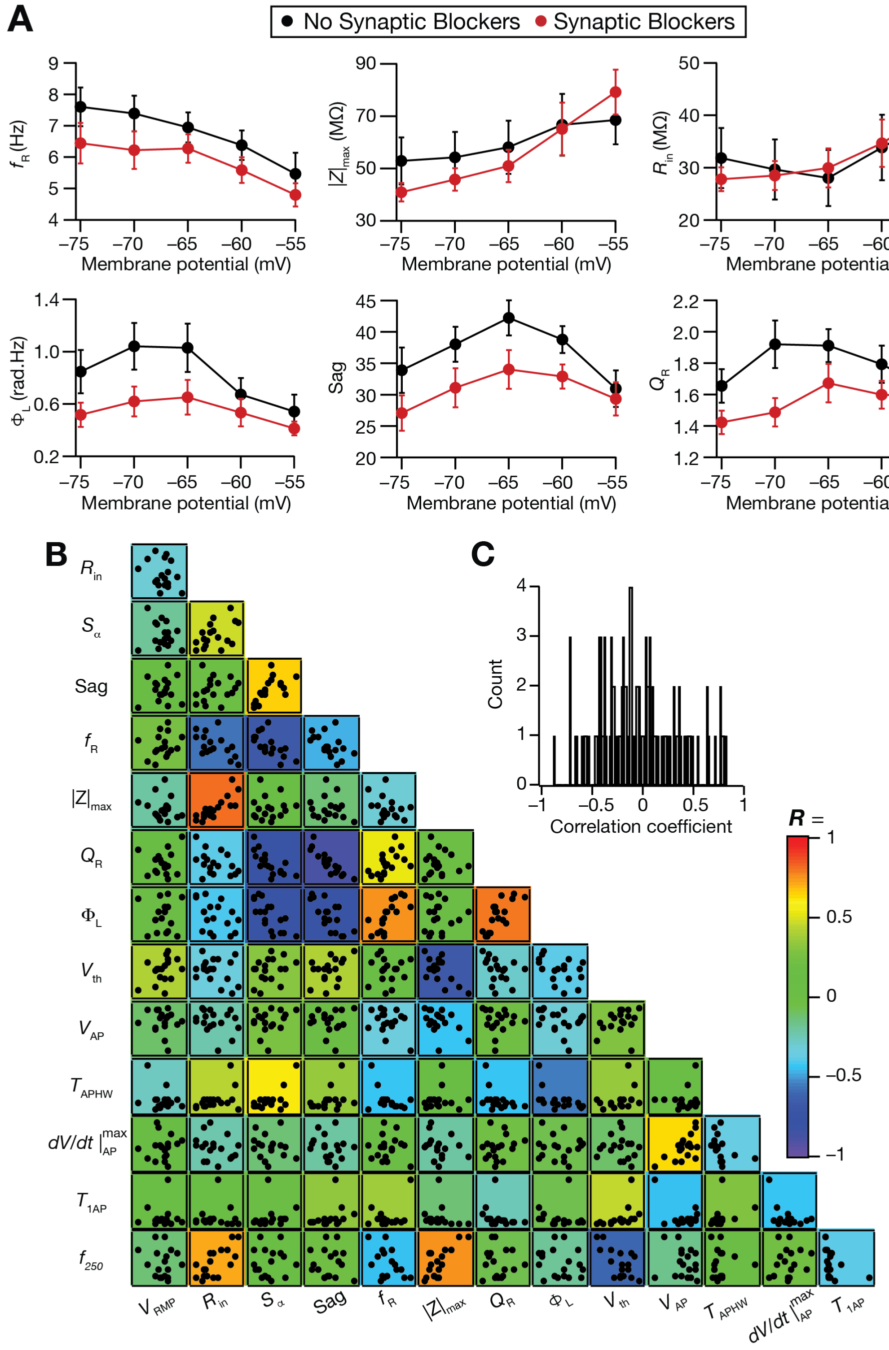

**Supplementary Figure S3: Voltage-dependent properties of sub-threshold measurements and pairwise correlations across sub and supra-threshold measurements for electrophysiologically recorded LII MEC stellate cells.** (A) Voltage dependence of sub-threshold response properties of stellate cells recorded with (red) or without (black) synaptic receptor blockers. Six sub-threshold measurements for each recorded cell were recorded by holding the cell at different voltages spanning –75 mV to –55 mV in steps of 5 mV. (B) Lower diagonal matrix depicting the pairwise scatter plot between 14 sub- and supra-threshold measurements from 28 LII MEC stellate cells (13 cells with and 15 without synaptic blockers). Individual scatterplots are overlaid on a heat map that depicts the pairwise correlation coefficient computed for that scatter plot. Inset: distribution of the correlation coefficient values from scatter plots in B.

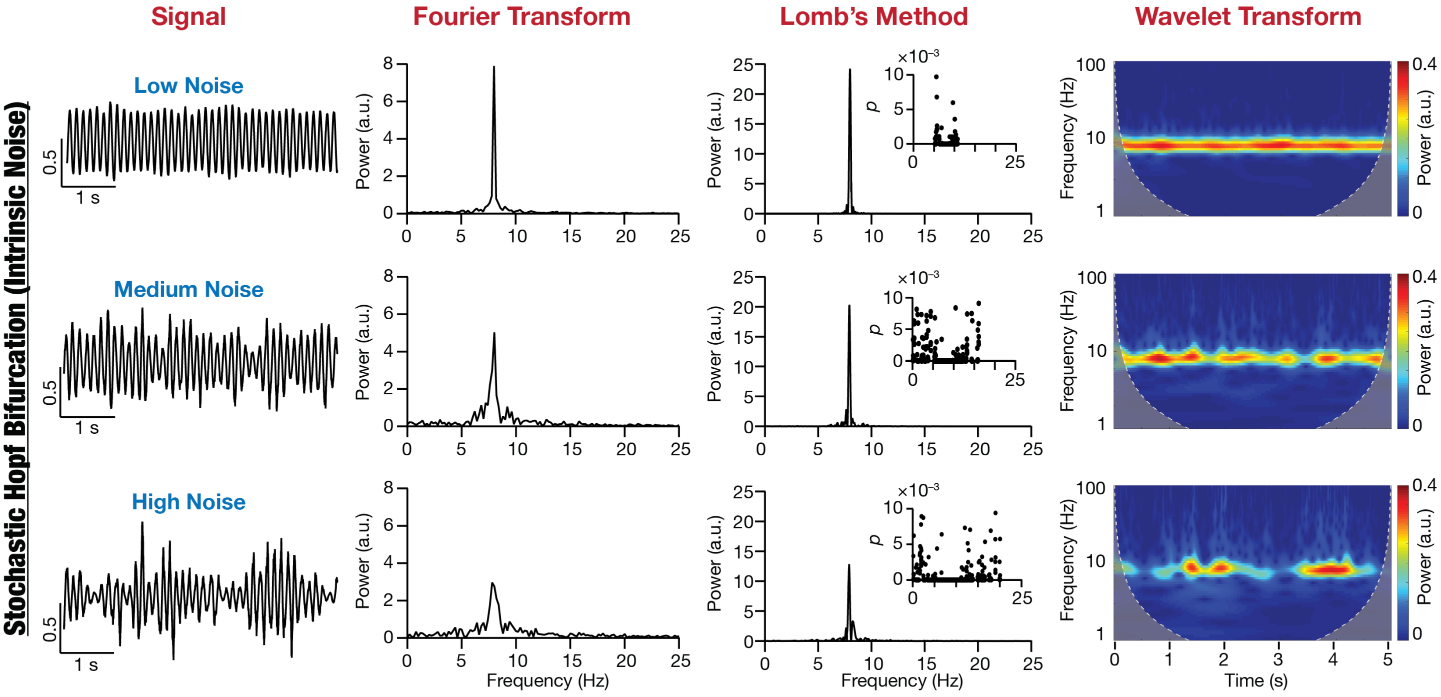

**Supplementary Figure S4: Spectral properties of oscillations emergent from a stochastic non-linear dynamical system with intrinsic noise**. Impact of parametric (intrinsic) noise on oscillations emerging from a nonlinear dynamical system (Hopf bifurcation). *Row 1*: Low noise, *Row 2*: Medium noise, and *Row 3*: High noise. *Column 1*: time-domain signal, *Column 2*: Fourier Transform (notice the 8 Hz peak) of the signal, *Column 3*: Lomb's periodogram of the signal with inset depicting the significance of each peak in the periodogram, and *Column 4*: spectrogram of the signal computed using wavelet transform.

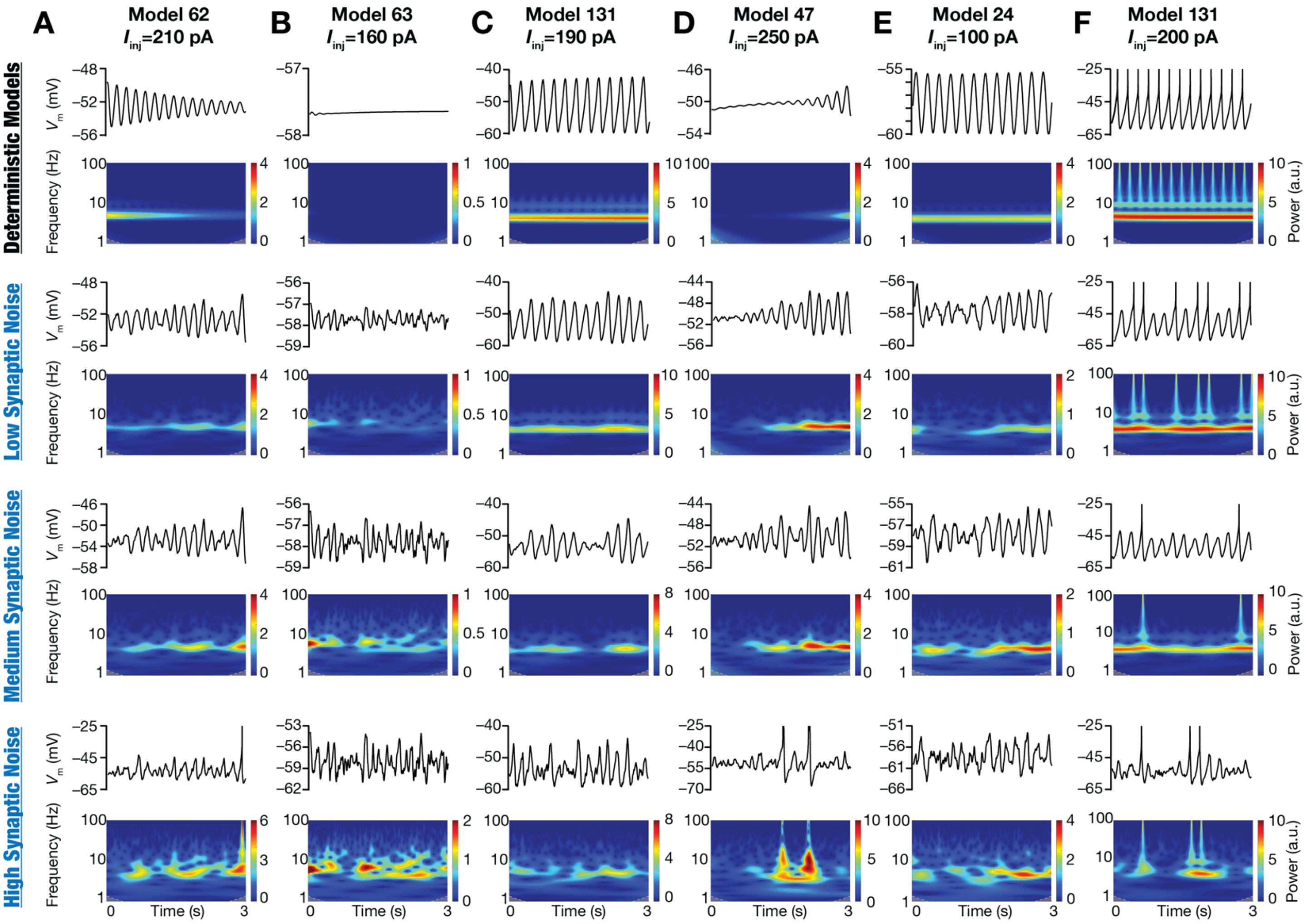

**Supplementary Figure S5: Illustrative examples of the role of synaptic noise in stabilizing peri-threshold oscillatory patterns in a heterogeneous population of LII MEC stellate cell models.** Each row in panels (A–F) depicts intrinsic activity patterns from different models for a 3 s period and the corresponding spectrograms computed using wavelet transform. Each column is identified by the corresponding model number along with identified values of injected current ($I_{\text{inj}}$) employed to generate the activity patterns. The first row depicts activity patterns in the deterministic model, in the absence of any form of noise. Rows 2–4 depict activity patterns from the same model, injected with the same $I_{\text{inj}}$ value, with low, medium, and high levels of synaptic noise. Note when spikes occurred, they were truncated to –25 mV to emphasize subthreshold dynamics.

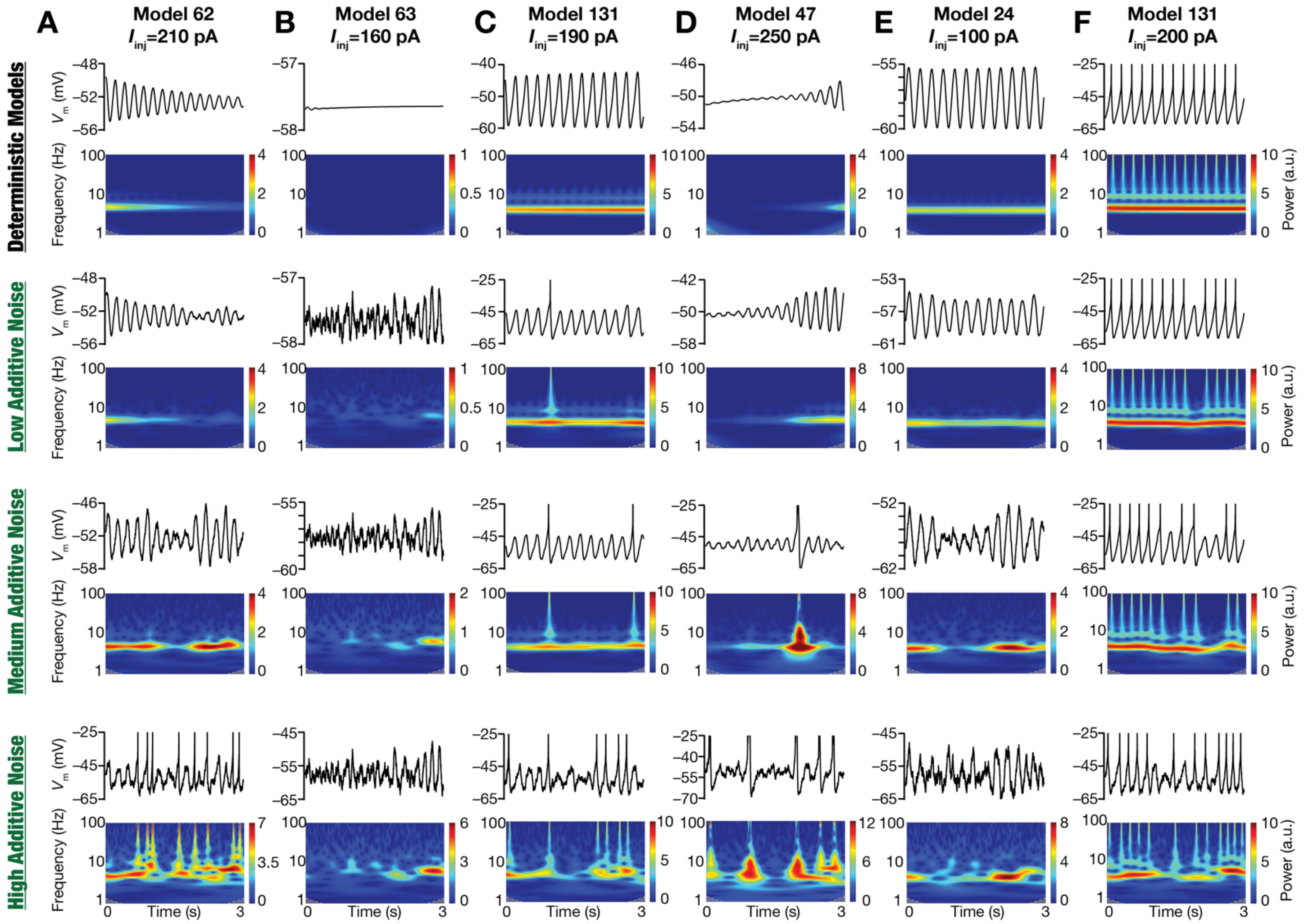

**Supplementary Figure S6: Illustrative examples of the role of external additive noise in stabilizing peri-threshold oscillatory patterns in a heterogeneous population of LII MEC stellate cell models.** Each row in panels (A–F) depicts intrinsic activity patterns from different models for a 3 s period and the corresponding spectrograms computed using wavelet transform. Each column is identified by the corresponding model number along with identified values of injected current ($I_{\text{inj}}$) employed to generate the activity patterns. The first row depicts activity patterns in the deterministic model, in the absence of any form of noise. Rows 2–4 depict activity patterns from the same model, injected with the same $I_{\text{inj}}$ value, with low (0.03), medium (0.12), and high (0.48) levels of additive noise. Note when spikes occurred, they were truncated to –25 mV to emphasize subthreshold dynamics.

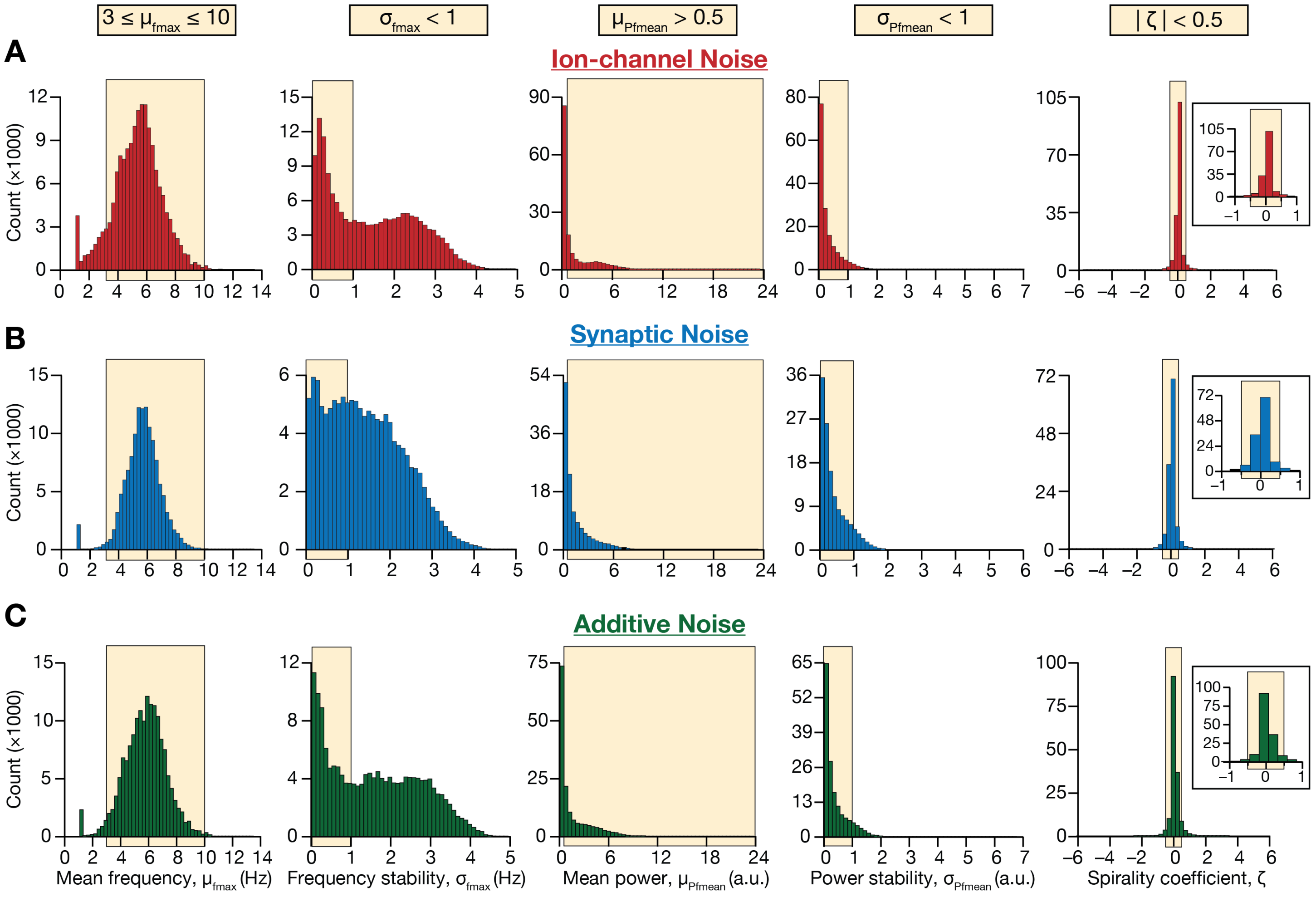

**Supplementary Figure S7: Distributions of spectrogram-based quantitative metrics used for assessing robustness in theta-frequency oscillatory activity in a stochastic, heterogeneous population of entorhinal stellate cells.** Distribution of the five spectrogram-based quantitative metrics for assessing robustness in theta-frequency oscillatory activity. *Column 1*, Mean frequency at maximal power, $\mu_{\text{fmax}}$; *Column 2*, Standard deviation of frequency at maximal power, $\sigma_{\text{fmax}}$; *Column 3*, Mean power at mean frequency, $\mu_{\text{Pfmean}}$; *Column 4*, Standard deviation of power at mean frequency, $\sigma_{\text{Pfmean}}$; *Column 5*, Spirality coefficient, $\zeta$. This population of measurements were derived from all model neurons ($n$ = 155), spanning all 21 current injection ($I_{\text{inj}}$) values, 10 independent trials for each level of the three forms of noise: (A) ion-channel noise, (B) synaptic noise, or (C) additive noise. The beige rectangle in each graph represents the bounds on the respective measurements for detection of a valid oscillatory trace.

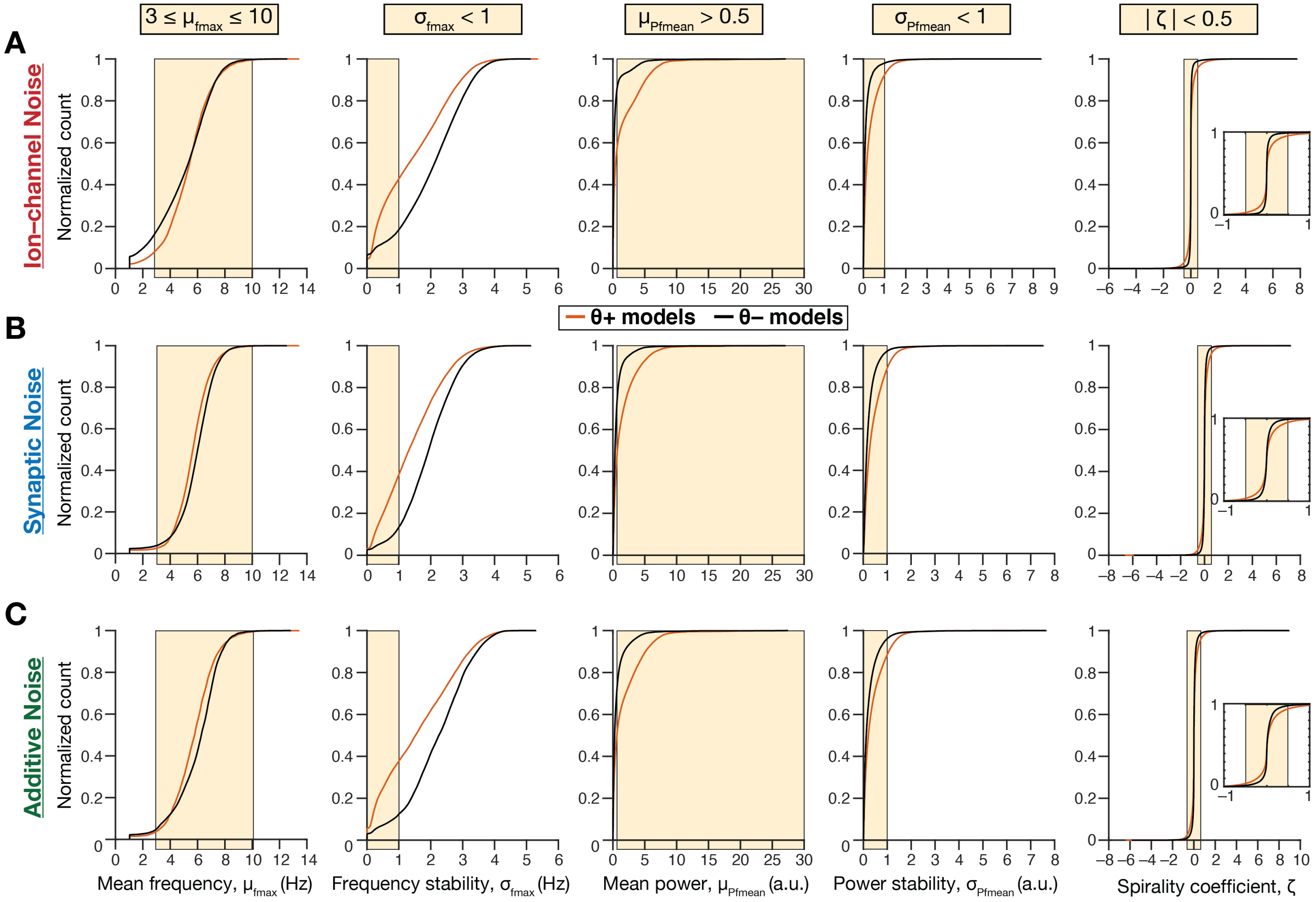

**Supplementary Figure 8: Cumulative distributions of spectrogram-based quantitative metrics used for assessing robustness in theta-frequency oscillatory activity in θ+ and θ– model cell populations.** Cumulative distributions of the five spectrogram-based quantitative metrics for assessing robustness in theta-frequency oscillatory activity in θ+ and θ– model cell populations. *Column 1*, Mean frequency at maximal power, $\mu_{\mathrm{fmax}}$; *Column 2*, Standard deviation of frequency at maximal power, $\sigma_{\mathrm{fmax}}$; *Column 3*, Mean power at mean frequency, $\mu_{\mathrm{Pfmean}}$; *Column 4*, Standard deviation of power at mean frequency, $\sigma_{\mathrm{Pfmean}}$; *Column 5*, Spirality coefficient, ζ. These populations of measurements were derived from all all θ+ ($n_{\theta+}$ = 155) and θ– ($n_{\theta-}$ = 155), spanning all 21 current injection ($I_{\mathrm{inj}}$) values, 10 independent trials for each level of the three forms of noise: (A) ion-channel noise, (B) synaptic noise, or (C) additive noise. The beige rectangle in each graph represents the bounds on the respective measurements for detection of a valid oscillatory trace.

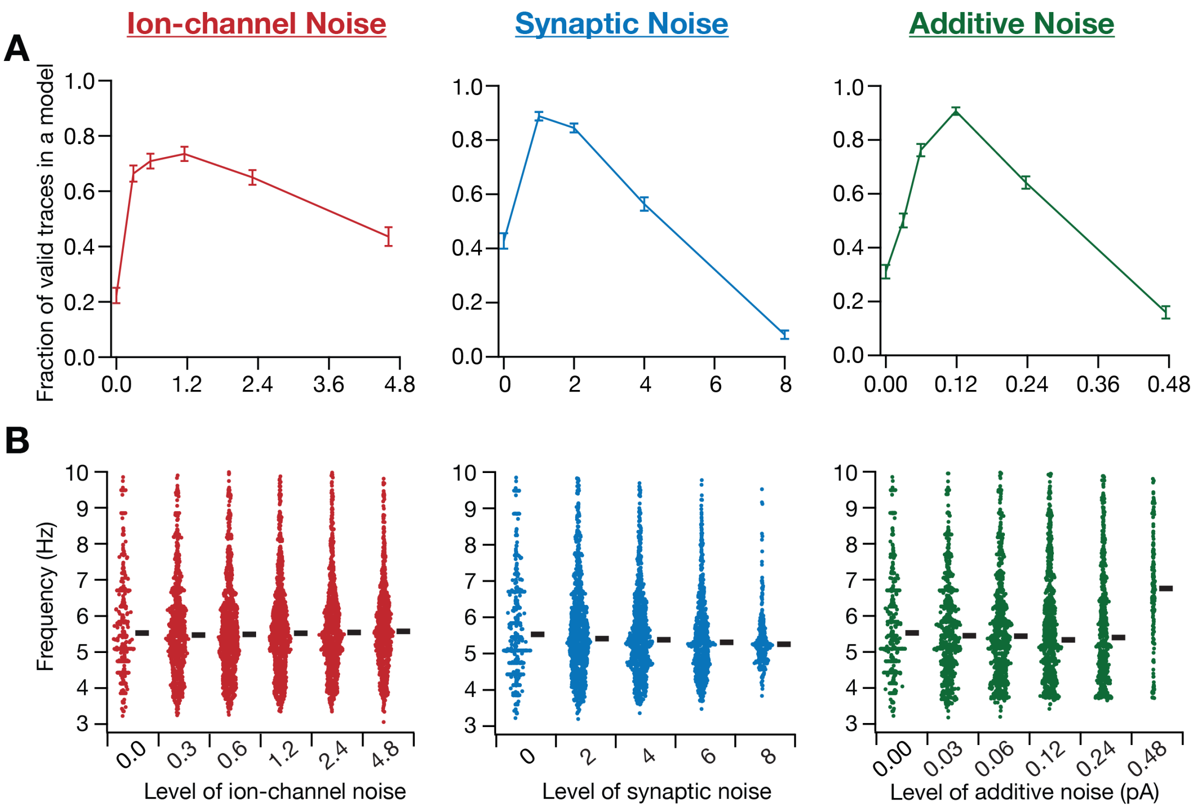

**Supplementary Figure 9: Stochastic resonance in the emergence of peri-threshold theta-frequency oscillations at the single-neuron level.** (A) The fraction of valid oscillatory traces, normalized for each neuron across noise levels, plotted (mean ± SEM) as a function of level of each form of noise. (B) Oscillatory frequency of all valid oscillatory traces spanning different levels of each form of noise. The thick black line beside each plot represents the median for those set of frequencies.

**Supplementary Table S1: Measurement statistics from stellate cells in the absence of synaptic blockers.**

| Measurement | Mean | SEM | SD | CV | Median | IQR |
|---|---|---|---|---|---|---|
| $V_{\mathrm{RMP}}$ (mV) | –58.79 | 0.76 | 2.93 | –0.05 | –58.79 | 2.89 |
| $S_{\alpha}$ | 0.79 | 0.04 | 0.14 | 0.18 | 0.74 | 0.23 |
| $R_{\mathrm{in}}$ (MΩ) | 42.02 | 4.68 | 18.11 | 0.43 | 41.44 | 32.86 |
| *Sag* | 0.62 | 0.02 | 0.09 | 0.15 | 0.60 | 0.11 |
| $f_{\mathrm{R}}$ (Hz) | 6.10 | 0.42 | 1.64 | 0.27 | 5.91 | 2.26 |
| $\lvert Z\rvert_{\mathrm{max}}$ (MΩ) | 74.54 | 9.74 | 37.73 | 0.51 | 67.32 | 38.33 |
| $Q_{\mathrm{R}}$ | 1.79 | 0.11 | 0.43 | 0.24 | 1.79 | 0.66 |
| $\Phi_{\mathrm{L}}$ (rad.Hz) | 0.58 | 0.12 | 0.45 | 0.77 | 0.41 | 0.60 |
| $V_{\mathrm{th}}$ (mV) | –43.84 | 1.75 | 5.26 | –0.12 | –42.43 | 4.67 |
| $V_{\mathrm{AP}}$ (mV) | 103.61 | 1.56 | 4.69 | 0.05 | 105.07 | 5.34 |
| *SFA* | 0.39 | 0.07 | 0.19 | 0.49 | 0.39 | 0.28 |
| $T_{1\mathrm{ISI}}$ (ms) | 28.84 | 8.83 | 26.49 | 0.92 | 27.20 | 17.35 |
| $T_{\mathrm{APHW}}$ (ms) | 0.80 | 0.07 | 0.20 | 0.24 | 0.72 | 0.08 |
| $\left.\frac{dV}{dt}\right\rvert_{\mathrm{AP}}^{\mathrm{max}}$ (V/s) | 441.82 | 24.13 | 72.39 | 0.16 | 419.91 | 119.63 |

**Supplementary Table S2: Measurement statistics from stellate cells in the presence of synaptic blockers.**

| Measurement | Mean | SEM | SD | CV | Median | IQR |
|---|---|---|---|---|---|---|
| $V_{\mathrm{RMP}}$ (mV) | –58.87 | 0.94 | 3.38 | –0.06 | –58.21 | 3.33 |
| $S_{\alpha}$ | 0.82 | 0.02 | 0.06 | 0.07 | 0.81 | 0.07 |
| $R_{\mathrm{in}}$ (MΩ) | 38.68 | 3.63 | 13.09 | 0.34 | 37.18 | 14.38 |
| *Sag* | 0.69 | 0.02 | 0.08 | 0.11 | 0.71 | 0.09 |
| $f_{\mathrm{R}}$ (Hz) | 5.32 | 0.34 | 1.22 | 0.23 | 5.25 | 1.55 |
| $\lvert Z\rvert_{\mathrm{max}}$ (MΩ) | 70.38 | 7.17 | 25.84 | 0.37 | 61.03 | 29.94 |
| $Q_{\mathrm{R}}$ | 1.53 | 0.07 | 0.26 | 0.17 | 1.51 | 0.32 |
| $\Phi_{\mathrm{L}}$ (rad.Hz) | 0.40 | 0.06 | 0.20 | 0.51 | 0.40 | 0.31 |
| $V_{\mathrm{th}}$ (mV) | –41.74 | 1.63 | 4.88 | –0.12 | –41.01 | 7.40 |
| $V_{\mathrm{AP}}$ (mV) | 100.30 | 1.85 | 5.56 | 0.06 | 100.62 | 4.20 |
| *SFA* | 0.36 | 0.10 | 0.21 | 0.60 | 0.23 | 0.24 |
| $T_{1\mathrm{ISI}}$ (ms) | 49.63 | 26.19 | 78.57 | 1.58 | 22.42 | 19.34 |
| $T_{\mathrm{APHW}}$ (ms) | 0.71 | 0.02 | 0.06 | 0.08 | 0.70 | 0.10 |
| $\left.\frac{dV}{dt}\right\rvert_{\mathrm{AP}}^{\mathrm{max}}$ (V/s) | 394.21 | 16.69 | 50.06 | 0.13 | 421.13 | 92.16 |

**SEM**: Standard error of the mean; **SD**: Standard deviation; **CV**: Coefficient of variation; **IQR**: Interquartile range.
